# A selective projection from the subthalamic nucleus to parvalbumin-expressing interneurons of the striatum

**DOI:** 10.1101/2020.11.26.400242

**Authors:** Krishnakanth Kondabolu, Natalie M. Doig, Olaoluwa Ayeko, Bakhtawer Khan, Alexandra Torres, Daniela Calvigioni, Konstantinos Meletis, Peter J. Magill, Tibor Koós

## Abstract

The striatum and subthalamic nucleus (STN) are considered to be the primary input nuclei of the basal ganglia. Projection neurons of both striatum and STN can extensively interact with other basal ganglia nuclei, and there is growing anatomical evidence of direct axonal connections from the STN to striatum. There remains, however, a pressing need to elucidate the organization and impact of these subthalamostriatal projections in the context of the diverse cell types constituting the striatum. To address this, we carried out monosynaptic retrograde tracing from genetically-defined populations of dorsal striatal neurons in adult male and female mice, quantifying the connectivity from STN neurons to spiny projection neurons, GABAergic interneurons, and cholinergic interneurons. In parallel, we used a combination of *ex vivo* electrophysiology and optogenetics to characterize the responses of a complementary range of dorsal striatal neuron types to activation of STN axons. Our tracing studies showed that the connectivity from STN neurons to striatal parvalbumin-expressing interneurons is significantly higher (~ four-to eight-fold) than that from STN to any of the four other striatal cell types examined. In agreement, our recording experiments showed that parvalbumin-expressing interneurons, but not the other cell types tested, commonly exhibited robust monosynaptic excitatory responses to subthalamostriatal inputs. Taken together, our data collectively demonstrate that the subthalamostriatal projection is highly selective for target cell type. We conclude that glutamatergic STN neurons are positioned to directly and powerfully influence striatal activity dynamics by virtue of their enriched innervation of GABAergic parvalbumin-expressing interneurons.

## Introduction

Influential models of the functional organization of basal ganglia (BG) circuits and their thalamic and cortical partners, such as the ‘direct/indirect pathways’ scheme (DeLong, 1990; Smith et al., 1998) and a modification accentuating the ‘hyperdirect pathway’ (Nambu et al., 2002), consider the striatum and subthalamic nucleus (STN) to be the primary input nuclei of the BG. The same models assume that striatum and STN are not monosynaptically connected. Accordingly, the primary targets of striatal and STN projecting axons are typically listed as the external globus pallidus (GPe) and BG output nuclei *i.e.* internal globus pallidus (or entopeduncular nucleus in rodents) and substantia nigra pars reticulata (Smith et al., 1998; Emmi et al., 2020). That striatum is not commonly deemed a target of STN outputs is at variance with anatomical evidence. Indeed, studies of neuronal populations using ‘classical’ tracers support the existence of moderate subthalamostriatal projections in rats, cats and monkeys (Beckstead, 1983; Kita and Kitai, 1987; Nakano et al., 1990; Smith et al., 1990), although data can be obfuscated by the bimodal (mixed anterograde and retrograde) transport of these tracers and their potential uptake by fibers of passage (Smith et al., 1998). On the other hand, single-neuron tracing indicates that most rat STN neurons innervate striatum, and sometimes to a greater degree than innervation of BG output nuclei (Koshimizu et al., 2013). Furthermore, recent studies employing sophisticated viral vector-mediated cell labeling in mice suggest that STN neuron axons can form synapses with striatal neurons (Wall et al., 2013; Guo et al., 2015; Smith et al., 2016; Klug et al., 2018; Choi et al., 2019). Importantly, there has been no physiological characterization of the incidence and strength of subthalamostriatal neurotransmission.

Understanding the organization and impact of the subthalamostriatal projection is challenged by the diversity of cell types within striatum. Approximately 90-95% of striatal neurons can be classified as spiny projection neurons (SPNs), which may then be further divided into, for example, populations giving rise to the direct pathway or indirect pathway (Gerfen and Surmeier, 2011). The remaining ~5-10% is made up of interneurons that are either cholinergic or GABAergic, the latter of which also exhibit substantial heterogeneity in form and function (Tepper et al., 2010, 2018). Stemming from this diversity, there is much scope for axons to engage one or more cell types in a biased manner. Indeed, the connections between striatal neurons are often selective for cell type, supporting a complex mix of reciprocal and non-reciprocal interactions across the wider microcircuit. Some of the extrinsic inputs to striatum, arising from both cortical and subcortical areas, also exhibit selectivity for target cells (Silberberg and Bolam, 2015; Tepper et al., 2018; Assous and Tepper, 2019; Assous et al., 2019).

Currently, it is unclear how the STN innervates and influences the striatal microcircuit, including the extent to which the subthalamostriatal projection is selective for target neuron type(s). Resolving these issues requires definitions of the structural substrates and physiological properties of this projection, while accounting for cellular diversity in striatum. To address this, we carried out monosynaptic retrograde tracing from distinct populations of striatal neurons in genetically-altered mice, quantifying the connectivity from STN neurons to SPNs, GABAergic interneurons and cholinergic interneurons. To complement these anatomical studies and gain insights into neurotransmission, we used *ex vivo* electrophysiology and optogenetics to interrogate the responses of a corresponding range of striatal cell types to activation of subthalamostriatal axons. Our results collectively reveal that the subthalamostriatal projection is highly selective, providing relatively rich and efficacious glutamatergic inputs to GABAergic parvalbumin-expressing interneurons.

## Materials and Methods

Experimental procedures were performed on mice and were conducted either at the University of Oxford in accordance with the Animals (Scientific Procedures) Act, 1986 (United Kingdom), or at Rutgers University with the approval of the Institutional Animal Care and Use Committee (IACUC) in accordance with public health service (PHS) policy on humane care and use of laboratory animals.

### Animals and related procedures for anterograde labeling and monosynaptic retrograde labeling of neurons in vivo

Adult male and female mice, aged 2.5-8 months, were used for these experiments. Eight lines of genetically-altered mice were used; all were bred to a C57Bl6/J background, and only mice heterozygous/hemizygous for the transgene(s) were used in experiments. To target glutamatergic neurons of the STN for anterograde labeling, we used a VGluT2-Cre mouse line (B6J.129S6(FVB)-*Slc17a6^tm2(cre)Lowl^*/MwarJ; Jackson Laboratory; RRID:IMSR_JAX:028863). To retrogradely label STN neurons innervating striatal neurons as a whole, we used a ‘double transgenic’ line produced by crossing VGAT-Cre mice (*Slc32a1^tm2(cre)Lowl^*/J; Jackson Laboratory; RRID:IMSR_JAX:016962) with ChAT-Cre mice (B6;129S6-*Chat^tm2(cre)Lowl^*/J; Jackson Laboratory; RRID:IMSR_JAX:006410). To retrogradely label STN neurons innervating more restricted populations of neurons in striatum, we used the following lines: Drd1a-Cre mice (B6.FVB(Cg)-Tg(Drd1-cre)EY262Gsat/Mmucd; GENSAT/MMRRC; RRID:MMRRC_030989-UCD); Adora2a-Cre mice (B6.FVB(Cg)-Tg(Adora2a-cre)KG139Gsat/Mmucd; GENSAT/MMRRC; RRID:MMRRC_036158-UCD); PV-Cre mice (B6;129P2-*Pvalb^tm1(cre)Arbr^*/J; Jackson Laboratory; RRID:IMSR_JAX:008069); SOM-Cre mice (*Sst^tm2.1(cre)Zjh^*/J; Jackson Laboratory; RRID:IMSR_JAX:013044); and ChAT-Cre mice (B6;129S6-*Chat^tm2(cre)Lowl^*/J; Jackson Laboratory; RRID:IMSR_JAX:006410).

For stereotaxic intracerebral injections of adeno-associated virus (AAV) and rhabdovirus vectors in mice, general anesthesia was induced and maintained with isoflurane (1.0-3.0% v/v in O_2_). Animals received perioperative analgesic (buprenorphine, 0.1 mg/kg, s.c.; Ceva) and were placed in a stereotaxic frame (Kopf Instruments). Wound margins were first infiltrated with local anesthetic (0.5% w/v bupivacaine [Marcaine]; Aspen). Body temperature was maintained at ~37°C by a homeothermic heating device (Harvard Apparatus). To anterogradely label glutamatergic STN neurons with green fluorescent protein (GFP), we used a glass micropipette (internal tip diameter of 15-22 μm) to unilaterally or bilaterally inject ~33 nl (per site) of a AAV2-CAG-FLEX-GFP vector (University of North Carolina [UNC] Vector Core) into the STN of VGluT2-Cre mice, using the following stereotaxic coordinates: 1.90 mm posterior of Bregma, 1.75 mm lateral of Bregma, and 4.75 mm ventral to the brain surface. To minimize reflux, the micropipette was left in place for ~20 min after the injection. Mice were maintained for 28-35 d after surgery to allow for neuron transduction and labeling with GFP; they were then humanely killed and perfused (see below). To carry out monosynaptic retrograde tracing from neurons in the dorsal striatum, we made sequential use of a single Cre-dependent ‘helper virus’ (AAV5-DIO-TVA^V5^-RG; bicistronically expressing TVA receptor fused to a V5 tag, and the rabies glycoprotein [RG]; (Ährlund-Richter et al., 2019)) and a ‘modified rabies virus’ ([EnvA]-SADΔG-EGFP; pseudotyped with EnvA, RG deleted, and expressing enhanced GFP; (Fürth et al., 2018; Ährlund-Richter et al., 2019)). In a first step, we used a glass micropipette to unilaterally inject 60-120 nl of helper virus into the central aspects of the dorsal striatum of Cre-expressing mice, using the following coordinates (caudal approach at an angle of 20° to vertical): 0.00 mm anterior of Bregma, 2.20 mm lateral of Bregma, and 2.70 mm ventral to the brain surface. To minimize reflux, the micropipette was left in place for ~10 min after the injection. Allowing 21 d for neuron transduction, Cre-mediated recombination, and the generation of ‘starter’ neurons expressing all components necessary for retrograde labeling of their presynaptic partners, we then as a second step injected 60-120 nl of modified rabies virus into the same striatal locations, using the following coordinates (vertical, no angle): 1.00 mm anterior of Bregma, 2.20 mm lateral of Bregma, and 2.50 mm ventral to the brain surface. To minimize reflux, the micropipette was left in place for ~10 min after the injection.

Mice were maintained for 7 d after surgery to allow for starter neuron infection and retrograde trans-synaptic labeling of neurons providing monosynaptic inputs to starters. Thereafter, mice were killed with pentobarbitone (1.5 g/kg, i.p.; Animalcare) and transcardially perfused with 20-50 ml of 0.05 M PBS, pH 7.4 (PBS), followed by 30-100 ml 4% w/v PFA in 0.1 M phosphate buffer, pH 7.4 (PB). Brains were removed and left overnight in fixative at 4°C before sectioning.

### Tissue processing, indirect immunofluorescence, imaging, and stereological quantification for anterograde/retrograde labeling experiments in vivo

Brains were embedded in agar (3-4% w/v dissolved in dH_2_O) before being cut into 50 μm coronal sections on a vibrating microtome (VT1000S; Leica Microsystems). Free-floating tissue sections were collected in series, washed in PBS, and stored in PBS containing 0.05% w/v sodium azide (Sigma) at 4°C until processing for indirect immunofluorescence to reveal molecular markers (Abdi et al., 2015). Briefly, after 1 h of incubation in “Triton PBS” (PBS with 0.3% v/v Triton X-100 and 0.02% w/v sodium azide) containing 10% v/v normal donkey serum (NDS; 017-000-121, Jackson ImmunoResearch Laboratories, RRID:AB_2337258), sections were incubated overnight at room temperature, or for 72 h at 4°C, in Triton PBS containing 1% v/v NDS and a mixture of between two and four of the following primary antibodies: goat anti-choline acetyltransferase (ChAT; 1:500 dilution; AB144P, Millipore, RRID:AB_2079751); goat anti-forkhead box protein P2 (FoxP2; 1:500; sc-21069, Santa Cruz Biotechnology, RRID:AB_2107124); rat anti-GFP (1:1000; 04404-84, Nacalai Tesque, RRID:AB_10013361); guinea pig anti-neuronal nuclei protein (NeuN, also known as hexaribonucleotide-binding protein 3; 1:500, 266004, Synaptic Systems, RRID:AB_2619988); goat anti-nitric oxide synthase (NOS; 1:500; ab1376; Abcam; RRID:AB_300614); guinea pig anti-parvalbumin (PV; 1:1000; 195004, Synaptic Systems, RRID:AB_2156476); Mouse anti-somatostatin (SOM; 1 in 250; GTX71935, GeneTex, RRID:AB_383280); chicken anti-V5 tag (V5; 1:2000; ab9113, Abcam, RRID:AB_307022); and rabbit anti-vesicular glutamate transporter 2 (VGluT2; 1:1000; 135403, Synaptic Systems, RRID:AB_887883). After exposure to primary antibodies, sections were washed in PBS and incubated overnight at room temperature in Triton PBS containing an appropriate mixture of secondary antibodies (all raised in donkey) with minimal cross-reactivity and that were conjugated to the following fluorophores: AMCA (1:250 dilution; Jackson ImmunoResearch Laboratories); Alexa Fluor 488 (1:1000; Invitrogen); Cy3 (1:1000; Jackson ImmunoResearch Laboratories), Cy5 or DyLight 647 (1:500; Jackson ImmunoResearch Laboratories). To optimize immunolabeling for COUP TF-interacting protein 2 (Ctip2, also known as Bcl11b) and preproenkephalin (PPE) in tissue sections from Drd1a-Cre and Adora2a-Cre mice, we used a heat pretreatment as a means of antigen retrieval (Mallet et al., 2012; Garas et al., 2016; Sharott et al., 2017). Following incubation of sections in primary antibodies to GFP (mouse anti-GFP 1:1000; A-11120, Invitrogen, RRID:AB_221568) and V5, and then secondary antibodies to reveal GFP and V5 (as above), sections were sequentially washed in PBS and citrate buffer (10 mM citric acid, pH 6) before incubation in citrate buffer at 80°C for 1 h. Sections were then allowed to come to room temperature, and washed back into PBS, before incubation overnight at room temperature in PBS containing 1% v/v NDS and primary antibodies to Ctip2 (rat anti-Ctip2; 1:500; ab18465, Abcam, RRID:AB_2064130) and PPE (rabbit anti-PPE; 1:5000; LS-C23084, LifeSpan, RRID:AB_902714). Sections were then washed in PBS and incubated overnight at room temperature in PBS containing an appropriate mixture of secondary antibodies (as above). For analyses of Ctip2 in tissue sections from VGAT-Cre:ChAT-Cre mice, we revealed immunoreactivity for GFP (guinea pig anti-GFP; 1:1000; 132005, Synaptic Systems, RRID:AB_11042617), V5 and ChAT (rabbit anti-ChAT; 1:1000; 297013, Synaptic Systems, RRID:AB_2620040) before heat pretreatment and revelation of immunoreactivity for Ctip2. After binding of primary and secondary antibodies, and final washing in PBS, sections were mounted on glass slides and cover-slipped using Vectashield Mounting Medium (H-1000, Vector Laboratories, RRID:AB_2336789) or SlowFade Diamond Antifade Mountant (S36972, ThermoFisher Scientific). Coverslips were sealed using nail varnish and slides stored at 4°C before imaging.

A version of design-based stereology, the ‘modified optical fractionator’, was used to generate unbiased cell counts and determine the proportions of a given population of neurons that expressed certain combinations of molecular markers (Abdi et al., 2015; Dodson et al., 2015; Garas et al., 2016). All stereology, imaging for stereology, and cell counting, was performed using Stereo Investigator software (v. 2019.1.4, MBF Bioscience, RRID:SCR_002526). Acquisition of tissue images for stereological sampling was carried out on an AxioImager.M2 microscope (Zeiss) equipped with an ORCA Flash-4.0 LT digital CMOS camera (Hamamatsu), an Apotome.2 (Zeiss), and a Colibri 7 LED light source (Type R[G/Y]B-UV, Zeiss). Appropriate sets of filter cubes were used to image the fluorescence channels: AMCA (excitation 299–392 nm, beamsplitter 395 nm, emission 420-470 nm); Alexa Fluor 488 (excitation 450–490 nm, beamsplitter 495 nm, emission 500-550 nm); Cy3 (excitation 532-558 nm, beamsplitter 570 nm, emission 570-640 nm); and Cy5/DyLight 649 (excitation 625-655 nm, beamsplitter 660 nm, emission 665-715 nm). Images of each of the channels were taken sequentially and separately to negate possible crosstalk of signal across channels. In order to quantify retrogradely-labeled subthalamostriatal neurons, we first defined (using a 10× objective lens; 0.45 NA; Plan-Apochromat, Zeiss) the borders of the STN according to the expression of FoxP2 (Abdi et al., 2015); the full extent of the STN was imaged for each series examined in each mouse. To quantify striatal starter neurons, we first defined (using a 10× objective) the outer boundaries of regions within striatum that contained neurons immunoreactive for V5, an indicator of neurons transduced with the helper virus (and thus, potential starter cells). After delineating the borders of STN and the boundaries of striatal regions containing starter neurons, images for stereological sampling were acquired using the optical fractionator workflow in Stereo Investigator, employing a 2 μm-thick ‘guard zone’, and an unbiased 2D counting frame and grid frame of 600 × 600 μm (i.e., 100% of the region in the X, Y plane was sampled). Z-stacked images across a 10 μm-thick ‘optical disector’ were acquired using a 20× objective lens (0.8 NA; Plan-Apochromat), and images or ‘optical sections’ were taken in 1 μm steps to ensure no loss of signal in the Z-axis. Captured images were then analyzed offline. A neuron was only counted once through the series of optical sections when its nucleus came into sharp focus within the disector; neurons with nuclei already in focus in the top optical section of the disector were ignored. The use of stereology, and this optical disector probe in particular, ensured that we could generate robust and unbiased cell counts in a timely manner. For a given molecular marker, X, we designate positive immunoreactivity (confirmed expression) as X+, and undetectable immunoreactivity (no expression) as X-. A neuron was classified as not expressing the tested molecular marker only when positive immunoreactivity could be observed in other cells on the same optical section as the tested neuron. Striatal starter neurons were defined by their co-expression of immunoreactivity for V5 and rabies-encoded GFP. To ensure a high level of precision in the cell counts, data were only included from individual mice when the Coefficient of Error (CE; using the Gundersen method) was ≤ 0.1 with a smoothness factor of m = 1 (West et al., 1991; Gundersen et al., 1999). The CE provides an estimate of sampling precision, which is independent of biological variance. As the value approaches zero, the uncertainty in the estimate precision reduces. The number of sections counted per mouse was thus dependent on variability; sections/series were added to the analysis until the CE ≤ 0.1 for each animal. Images for figures were acquired with a confocal microscope (LSM880, Zeiss). All image adjustments were linear and applied to every pixel.

### Preparation of animals for electrophysiological recordings and optogenetic manipulations of neurons ex vivo

Adult male and female mice, aged 3-10 months, were used for these experiments. Wildtype mice (C57Bl6/J) and three lines of genetically-altered mice were used; all genetically-altered mice were bred to a C57Bl6/J background, and only mice heterozygous/hemizygous for the transgene(s) were used in experiments. To visualize specified types of interneuron in striatum, we used the following lines: PV-tdTomato reporter mice (C57BL/6-Tg(*Pvalb-tdTomato*)^15Gfng^/J; Jackson Laboratory; RRID:IMSR_JAX:027395); NPY-GFP reporter mice (B6.FVB-Tg(*Npy-hrGFP)^1Lowl^*/J; Jackson Laboratory; RRID:IMSR_JAX:006417); and ChAT-Cre mice (B6; 129S6-*Chat^tm2(cre)Lowl^*/J; Jackson Laboratory; RRID:IMSR_JAX:006410).

For stereotaxic intracerebral injections of AAV vectors in mice, general anesthesia was induced and maintained with isoflurane (1.0-3.0% v/v in O_2_). Animals were placed in a stereotaxic frame (Kopf Instruments). Wound margins were first infiltrated with local anesthetic (0.25% w/v bupivacaine, with 1:200,000 epinephrine [Sensorcaine], Hospira). Body temperature was maintained at ~37°C by a homeothermic heating device. To express channelrhodopsin2 (ChR2) in STN neurons, we used a 1 μl microsyringe (7000 Series, Hamilton) to unilaterally inject ~45 nl of an AAV vector (all from UNC Vector Core) into the STN of mice, using the following stereotaxic coordinates: 1.88 mm posterior of Bregma, 1.68 mm lateral of Bregma, and 4.50 mm ventral to the brain surface. To minimize reflux, the microsyringe needle was left in place for ~10 min after the injection. For experiments using wildtype or PV-tdTomato mice, we injected an AAV5-CAMKIIa-hChR2(H134R)-EYFP vector into the STN. For experiments using NPY-GFP mice, we injected an AAV5-CAMKIIa-hChR2(H134R)-mCherry vector into the STN. For experiments using ChAT-Cre mice, we injected an AAV2-hSyn-hChR2(H134R)-EYFP vector into the STN. To visualize striatal cholinergic interneurons in the same ChAT-Cre mice, we unilaterally injected (ipsilateral to transduced STN) 600 nl of an AAV5-CAG-FLEX-tdTomato vector (UNC Vector Core) into the dorsal striatum, using the following stereotaxic coordinates (200 nl per site at three depths along the dorsal-ventral axis): 0.70 mm anterior of Bregma, 1.80 mm lateral of Bregma, and 3.20, 2.60 and 2.20 mm ventral to the brain surface. To minimize reflux, the microsyringe needle was left in place for ~10 min after the most dorsal injection. Animals received postoperative analgesic (buprenorphine SR-LAB, 0.1 mg/kg, s.c.; ZooPharm) and were maintained for 42-56 d after surgery to allow for neuron transduction; the mice were then deeply anesthetized and perfused for *ex vivo* recordings and anatomical verification of ChR2 expression (see below).

### Acute brain slice preparation, electrophysiological recordings and optogenetic manipulations of neurons ex vivo

Visualized recordings of striatal neurons were performed in brain slices acutely prepared from mice, as previously described (Faust et al., 2016; Assous et al., 2017). Briefly, mice were anesthetized with 3% v/v isoflurane in O_2_, followed by ketamine (100 mg/kg, i.p.; Henry Schien), and transcardially perfused with an ice-cold oxygenated *N*-methyl *D*-glucamine (NMDG)-based solution that contained the following (in mM): 103.0 NMDG, 2.5 KCl, 1.2 NaH_2_PO_4_, 30.0 NaHCO_3_, 20.0 HEPES, 25.0 dextrose, 101.0 HCl, 10.0 MgSO_4_, 2.0 Thiourea, 3.0 sodium pyruvate, 12.0 *N*-acetyl cysteine, 0.5 CaCl_2_ (saturated with 95% O_2_ and 5% CO_2_; 300-310 mOsm, pH 7.2-7.4). The brain was then quickly removed from the skull, blocked in the coronal or parasagittal plane, glued to the stage of a vibrating microtome (VT1200S; Leica Microsystems), and submerged in oxygenated ice-cold NMDG-based solution. Slices containing the dorsal striatum (250- or 300-μm thick) were then cut and transferred to a holding chamber containing oxygenated NMDG-based solution at 35°C for 5 min, after which they were transferred to another chamber containing oxygenated “artificial CSF” (ACSF) at 25°C and allowed to recover for at least 1 h before recording. The ACSF contained the following (in mM): 124 NaCl, 26 NaHCO_3_, 2.5 KCl, 1.2 NaH_2_PO_4_, 1 MgCl_2_, 2 CaCl_2_, 10 glucose, and 3 sodium pyruvate. The osmolarity and pH of the ACSF were 300-310 mOsm and 7.2-7.4, respectively. Additional slices containing the STN (250- or 300-μm thick) were cut where necessary and immersed in fixative (4% w/v PFA in PB for 1-2 d at 4°C) for *post hoc* anatomical verification of ChR2 expression in and around the STN (see below).

For electrophysiological testing, single slices of striatum were transferred to a recording chamber and perfused continuously with oxygenated ACSF maintained at 32-34°C. Slices were visualized using infrared gradient contrast video on a microscope (BX50WI, Olympus) equipped with a 40× water-immersion objective lens (LUMPIanFL/IR, Olympus), an iXon EMCCD camera (DU-885K-CS0, Andor), a halogen light source (TH3 power supply, Olympus), and imaging software (Solis v 4.4.0, Andor). A mercury lamp (BH2-RFL-T3 power supply, Olympus) coupled to appropriate filter cubes (U-M41001, Olympus; 49008, Chroma Technology) was used to visualize native fluorescence (of GFP, mCherry and/or tdTomato) in the slices, which in turn ensured accurate targeting of desired cell types in areas of striatum that were traversed by axons expressing fluorescent fusion proteins of ChR2. Somatic patch-clamp recordings of some individual striatal neurons were made in a whole-cell configuration (in voltage-clamp mode at a holding voltage (V_h_) of −70 mV, and/or in current-clamp mode) using glass pipettes that were filled with a K-gluconate-based solution composed of (in mM): 130 K-gluconate, 10 KCl, 2 MgCl_2_, 10 HEPES, 4 Na_2_ATP, 0.4 Na_3_GTP. Either 0.2% w/v biocytin (B4261, Sigma) or 1.5 μl/ml Alexa Fluor 594 dye (A10438, ThermoFisher Scientific) was added to the pipette solution to facilitate, in particular, the *post hoc* anatomical identification of SPNs (see below). The osmolarity and pH of this K-gluconate pipette solution were 290-295 mOsm and 7.3, respectively. Whole-cell recordings from other striatal neurons were made in voltage-clamp mode (V_h_ −70 mV) using glass pipettes that were filled with a CsCl-based solution composed of (in mM): 125 CsCl, 2 MgCl_2_, 10 HEPES, 4 Na_2_ATP, 0.4 Na_3_GTP, plus 1.5 μl/ml Alexa Fluor 594 dye. The osmolarity and pH of this CsCl pipette solution were 290-295 mOsm and 7.3, respectively. Irrespective of internal solution, pipettes typically exhibited a DC impedance of 3-5 MΩ measured in the recording chamber. Somatic patch-clamp recordings were obtained using a Multiclamp 700B amplifier (Molecular Devices, RRID:SCR_018455) and ITC-1600 digitizer (Instrutech), with AxoGraph software (v. 1.7.4, RRID:SCR_014284) used for data acquisition and analysis. Electrode signals were low-pass filtered (Bessel filter) at 1 kHz or 10 kHz for voltage-clamp or current-clamp recordings, respectively, and sampled at 20 kHz. Most SPNs were recorded in wildtype mice, and were classified as SPNs by their characteristic morphological properties and/or intrinsic electrophysiological properties (Gertler et al., 2008). When patched with pipettes containing the K-gluconate-based solution, SPNs were filled with either biocytin or Alexa Fluor 594 dye for *post hoc* anatomical analyses and verification of cell type (see below). Current-clamp recordings of spontaneous activity and responses to somatic current injections were also useful in confirming SPN identity. All SPNs patched with CsCl-based pipette solution were filled with Alexa Fluor dye for *post hoc* identification. Different types of striatal interneuron were targeted for recording according to their expression of native tdTomato or GFP fluorescence in slices from PV-tdTomato mice, NPY-GFP mice and ChAT-Cre mice.

To optically stimulate ChR2-expressing axons in striatum, brief flashes (2 or 5 ms) of blue light (470 nm, ~1.5 mW/mm^2^) were generated by a TTL-controlled LED (LB W5SN-GZJX-35-Z, Mouser Electronics; housed above the microscope condenser) and delivered to the slice as wide-field illumination. To avoid overt desensitization of ChR2, successive light flashes (single flash or, in some cases, 10-20 Hz trains of 5 flashes) were delivered at intervals of 15 s. We used two analytical approaches, based on averaging and latency, to assess whether optical stimuli evoked reliable monosynaptic responses in striatal neurons. Analysis of membrane dynamics, i.e. EPSPs or EPSCs, was performed on averaged responses from nine stimulus trials. Peak EPSP/EPSC amplitude was measured as the amplitude of the averaged event relative to averaged pre-stimulus baseline. Indicative amplitude thresholds for reliable detection of EPSPs and EPSCs were >0.3 mV and >2.5 pA, respectively. We considered evoked EPSPs/EPSCs to be putative monosynaptic responses when their onsets occurred at <6 ms from the onset of light flashes. We also analyzed a time window of 6-12 ms from flash onset to test for disynaptic/polysynaptic responses. We sometimes observed brief (<0.5 ms) ‘switching artifacts’ occurring within 0.1 ms of the onset and/or offset of light flashes; these artifacts were readily distinguished from evoked EPSPs/EPSCs. Series resistance was regularly monitored, but not compensated, during recordings; if there was >20% change in series resistance, neurons were excluded from further analyses. Theoretical liquid junction potential was estimated to be −10.3 mV, and was not corrected off-line in current-clamp recordings. After filling of neurons with biocytin or Alexa Fluor dye, slices were removed from the recording chamber and immersed in 4% w/v PFA in PB for 1-2 d at 4°C before further anatomical processing.

### Brain slice processing for histofluorescence and immunofluorescence

To verify recorded cell types and ChR2 expression in striatum, free-floating tissue slices (250-300 μm thick) were washed in Triton PBS and processed for histofluorescence and immunofluorescence. Biocytin-filled striatal neurons were revealed by incubating slices overnight at room temperature in Triton PBS containing Alexa Fluor 405-conjugated streptavidin (1:500 dilution; S32351, ThermoFisher Scientific). Axonal ChR2 expression was revealed by incubating slices for 1-2 h in Triton PBS containing 10% v/v NDS (D9663, Sigma, RRID:AB_2810235), then overnight at room temperature in Triton PBS containing 1% v/v NDS and Alexa Fluor 488-conjugated rabbit anti-GFP (1:500; A-21311, ThermoFisher Scientific, RRID:AB_221477). For *post hoc* verification of ChR2 expression in and around the STN, the expression of native enhanced yellow fluorescent protein (EYFP) fluorescence was compared to the borders of STN defined by FoxP2 immunolabeling: slices were incubated for 1-2 h in Triton PBS containing 10% v/v NDS, then overnight at room temperature in Triton PBS containing 1% v/v NDS and rabbit anti-FoxP2 (1:1000; HPA000382, Atlas Antibodies, RRID:AB_1078908), then washed in PBS, and then incubated at room temperature for 4-5 h in Triton PBS containing Alexa Fluor 594-conjugated donkey anti-rabbit IgG (1:1000; A-21207, Thermo Fisher Scientific, RRID:AB_141637). After binding of streptavidin/antibodies, and final washing in PBS, slices were mounted on glass slides, cover-slipped in Vectashield Mounting Medium, and imaged on a confocal microscope (FluoView FV1000, Olympus). The identities of recorded and biocytin-/dye-filled SPNs were verified according to the presence of densely spiny dendrites.

### Statistical analysis

For each experiment, descriptions of critical variables (e.g., number of mice, neurons, and other samples evaluated) as well as details of statistical design and testing can be found in the Results. For monosynaptic retrograde tracing experiments, the ratio (*x*/*y*) of the stereologically-estimated total number of striatal starter neurons (*x*) to the estimated total number of retrogradely-labeled STN neurons (*y*) was calculated for each animal (Do et al., 2016; Choi et al., 2019), and is referred to as the ‘normalized connectivity index’. Graphing and statistical analyses were performed with GraphPad Prism (v8.4.2 or v5.03, RRID:SCR_002798). The Shapiro-Wilk test was used to judge whether datasets were normally distributed (*p* ≤ 0.05 to reject). Significance for all statistical tests was set at *p* < 0.05 (exact *p* values are given in the text). Data are represented as group means ± SEMs unless stated otherwise, with some plots additionally showing individual samples (mice or neurons) as appropriate.

## Results

### Subthalamic nucleus neurons project to, and form synapses with, neurons in dorsal striatum

Evidence of a subthalamostriatal projection in mice is somewhat conflicting; one anterograde tracing study explicitly refutes its existence (Schweizer et al., 2016), whereas other retrograde tracing studies clearly support its existence (Tervo et al., 2016; Klug et al., 2018; Choi et al., 2019). We used a combination of anterograde and retrograde tracing strategies, actioned through intracerebral injections of ‘conditional’ viral vectors in transgenic mice, to provide further context on this issue and elucidate the structural basis for direct interactions between genetically-defined cell types in STN and striatum.

Most STN neurons in rodents and other mammalian species are glutamatergic and robustly express the *Slc17a6* gene encoding the synaptic protein vesicular glutamate transporter 2 (VGluT2) (Smith and Parent, 1988; Albin et al., 1989; Barroso-Chinea et al., 2007; Rico et al., 2010; Schweizer et al., 2016). To reveal the axonal projections of glutamatergic STN neurons, we stereotaxically injected AAV vectors expressing GFP in a Cre recombinase-dependent manner into the STN of adult *Slc17a6*-Cre (VGluT2-Cre) mice (Fig. 1*A*). The somata, dendrites and axons of transduced neurons were then revealed with immunoreactivity for GFP. This anterograde tracing strategy allowed for highly-selective transduction of neuronal somata and dendrites throughout the rostrocaudual extent of STN (Fig. 1*B-D*), delineated according to the borders of a group of densely-packed neurons immunoreactive for the transcription factor FoxP2 (Abdi et al., 2015). We occasionally observed sparse transduction of a few neuronal somata in the neighboring zona incerta and parasubthalamic nucleus (Fig. 1*B-D*). However, we did not observe any GFP-expressing (AAV-GFP+) somata in brain structures known to robustly innervate striatum, including the cerebral cortex, intralaminar and motor thalamus, basolateral and central amygdala, ventral tegmental area or substantial nigra *pars compacta* (Smith et al., 1998; Pan et al., 2010). The AAV-GFP+ axons of transduced STN neurons traversed rostrally into the dorsal striatum, where they diffusely arborized over several millimeters (Fig. 1*E-H*). The ventral striatum (nucleus accumbens) was also innervated by AAV-GFP+ axons, but the projection was comparatively scant (Fig. 1*E,F*). In contrast, we observed dense AAV-GFP+ axonal plexuses within the GPe (Fig. 1*H*), a major target of STN neurons (Smith et al., 1998). Within dorsal striatum, the AAV-GFP+ axons exhibited both ‘en passant’ and ‘aux terminaux’ boutons that were immunoreactive for VGluT2 (Fig. 1*I-L*). The results of these anterograde tracing studies support the existence of a glutamatergic projection from the STN to the dorsal striatum in mice.

**Figure 1.**
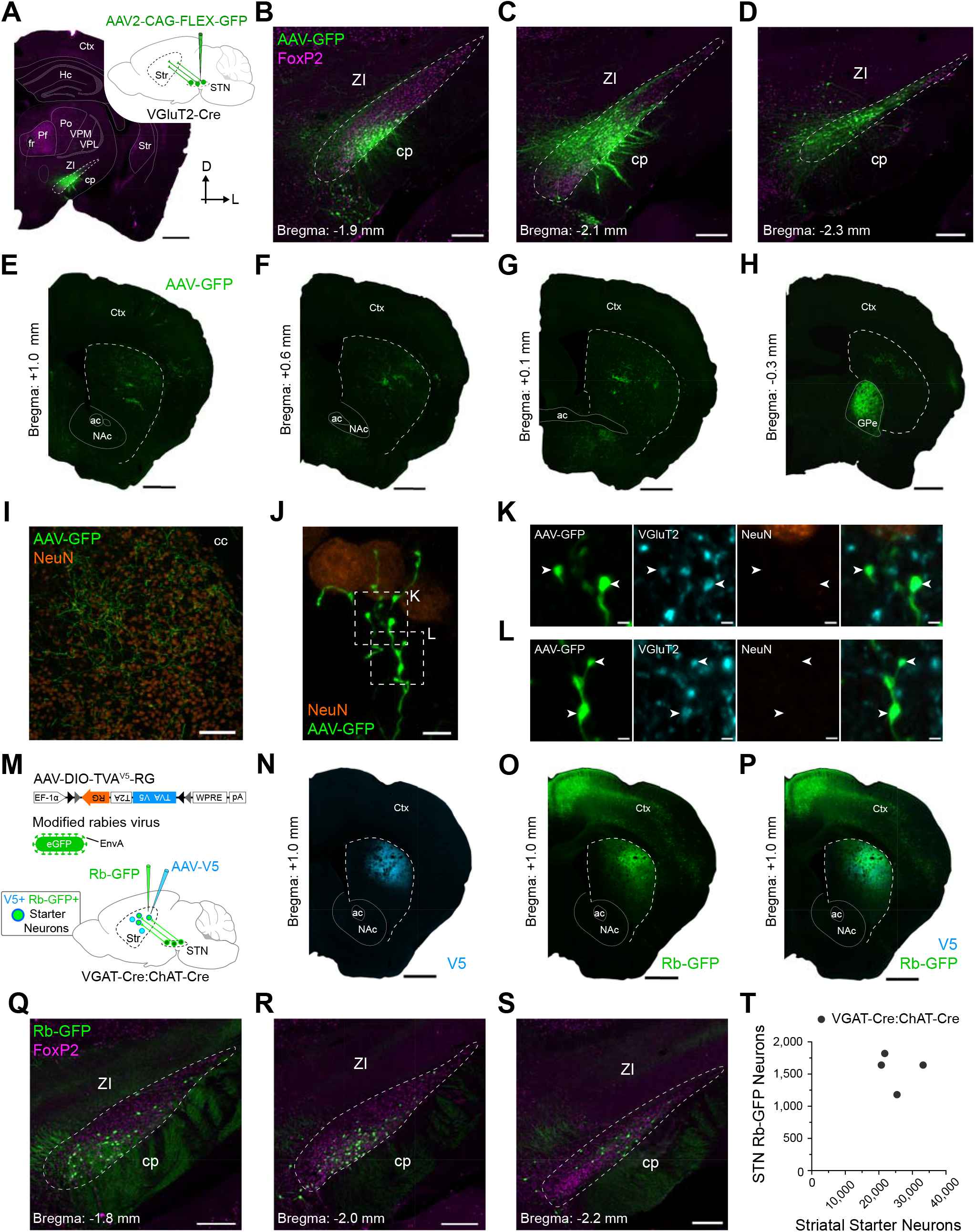
Subthalamic nucleus neurons can send axons to, and form synapses with, dorsal striatal neurons. ***A***, Strategy for selective anterograde tracing of glutamatergic subthalamic nucleus (STN) neurons with green fluorescent protein (GFP). *Inset,* a viral vector expressing GFP in a Cre recombinase-dependent manner (AAV2-CAG-FLEX-GFP) was unilaterally injected into the STN of adult VGluT2-Cre mice. Transduced anterogradely-labeled neurons were revealed with immunoreactivity for GFP (green). *Main image,* Coronal tissue section (D, dorsal; L, lateral) from a VGluT2-Cre mouse, showing a circumscribed group of transduced neurons in STN (boundaries marked by dashed lines). Immunoreactivity for the transcription factor FoxP2 (magenta) was used to facilitate delineation of STN and other brain regions. cp, cerebral peduncle; Ctx, cortex; fr, fascicularis retroflexus; Hc; hippocampus; Pf, parafascicular nucleus of the thalamus; Po, posterior thalamus; Str, striatum; VPL, ventral posterolateral nucleus; VPM, ventral posteromedial nucleus; ZI, zona incerta. ***B-D***, Immunofluorescence signals for anterogradely-labeled neurons (AAV-GFP) and FoxP2 in coronal sections from the same mouse as shown in ***A***. Note that AAV-GFP+ somata and dendrites were located in ‘rostral’ (***B***), ‘central’ (***C***), and ‘caudal’ (***D***) parts of STN. A standard stereotaxic reference (approximate distance relative to Bregma) is given for each rostrocaudal level. ***E-H***, Immunofluorescence signals for AAV-GFP+ axons in the dorsal striatum (boundaries marked by dashed lines) and other forebrain regions of the same mouse as that shown in ***A-D***. Sections are ordered from most rostral (***E***) to most caudal (***H***). Note the relatively dense plexus of AAV-GFP+ axons within the external globus pallidus (GPe), which is well known to be a major target of STN neurons (***H***). ac, anterior commissure; NAc, nucleus accumbens. ***I, J***, AAV-GFP+ axons exhibit boutons, both ‘en passant’ and ‘aux terminaux’, in dorsal striatum. Immunoreactivity for NeuN (orange) was used to reveal the somata of striatal neurons. ***K, L***, Higher-magnification confocal micrographs of boxed zones in ***J***. AAV-GFP+ axonal boutons expressed immunoreactivity for the vesicular glutamate transporter VGluT2 (cyan). ***M***, Strategy for retrograde labeling of STN neurons that monosynaptically innervate dorsal striatal neurons. A ‘helper virus’ (AAV-DIO-TVA^V5^-RG) and a ‘modified rabies virus’ (RG deleted, pseudotyped with EnvA, and expressing eGFP) were unilaterally and sequentially injected into the dorsal striatum of VGAT-Cre:ChAT-Cre mice. Striatal starter neurons, from which monosynaptic retrograde labeling of input neurons emanates, co-express V5 (blue) and rabies-encoded enhanced GFP (Rb-GFP, green). Retrogradely-labeled neurons in STN (and other brain regions) that innervate the starter neurons express Rb-GFP, but not V5. ***N-P***, Immunofluorescence signals for V5 (***N***), for Rb-GFP, as expressed by neurons transduced by the rabies virus (***O***), or for both V5 and Rb-GFP (***P***), in forebrain sections from a single VGAT-Cre:ChAT-Cre mouse. Note the enriched co-localization of V5 and Rb-GFP signals in the dorsal striatum. ***Q-S***, Retrogradely-labeled (Rb-GFP+) neurons were located in rostral (***Q***), central (***R***), and caudal (***S***) parts of the STN; all images from a single mouse. ***T***, Numbers of striatal starter neurons and retrogradely-labeled STN neurons in VGAT-Cre:ChAT-Cre mice *(n* = 4), as estimated using unbiased stereology (each purple circle indicates the estimates from a single mouse). Scale bars: ***A***, 750 μm; ***B-D***, 200 μm; ***E-H***, 750 μm; ***I***, 100 μm; ***J***, 5 μm; ***K, L***, 1 μm; ***N-P***, 750 μm; ***Q-S***, 200 μm.

To complement these anterograde labeling studies, we carried out monosynaptic retrograde tracing from neurons in the dorsal striatum (Callaway and Luo, 2015; Ährlund-Richter et al., 2019). This entailed the sequential injection of a Cre-dependent helper virus and a modified rabies virus into the dorsal striatum of Cre-expressing mice (Fig. 1*M*). We first aimed to elucidate the extent to which STN neurons innervate striatal neurons as a whole. The vast majority of striatal neurons are GABAergic, with the exception being cholinergic interneurons (Tepper et al., 2010, 2018; Silberberg and Bolam, 2015). These GABAergic neurons can be selectively accessed for study by exploiting their expression of the *Slc32a1* gene encoding the vesicular GABA transporter (VGAT), whereas cholinergic neurons can be targeted via their expression of the choline acetyltransferase (ChAT) gene. Thus, to achieve our aim, we unilaterally injected the Cre-dependent helper virus (bicistronically expressing TVA receptor fused to a V5 tag, and the rabies glycoprotein [RG]; (Ährlund-Richter et al., 2019)) into the central aspects of the dorsal striatum of double transgenic adult VGAT-Cre:ChAT-Cre mice (*n* = 4). Allowing time for Cre-mediated recombination, and the generation of ‘starter’ neurons expressing all components necessary for retrograde labeling of their presynaptic partners, we then injected modified rabies virus (RG deleted, pseudotyped with EnvA, and expressing enhanced GFP) into the same striatal locations (Fig. 1*M*). We visualized striatal starter neurons by their co-expression of immunoreactivity for V5 and rabies-encoded GFP (Rb-GFP; Fig. 1*M-P*). By virtue of stereotaxic targeting, we did not observe starter cells in the nucleus accumbens. The majority of starter neurons in dorsal striatum (86.9 ± 1.0% [mean ± S.E.M]; *n* = 5,055 total starter neurons counted from 4 mice) expressed nuclear immunoreactivity for Ctip2, a marker of most SPNs but not striatal interneurons (Arlotta et al., 2008). A small proportion of starter neurons (2.4 ± 0.4%) instead expressed somatic immunoreactivity for ChAT. A third group of starter neurons (10.7 ± 0.9%) expressed neither Ctip2 nor ChAT, suggesting they were GABAergic interneurons. Retrogradely-labeled neurons that monosynaptically innervated the starter neurons were identified by their expression of immunoreactivity for Rb-GFP, but not V5 (Fig. 1*M-P*). Importantly, Rb-GFP+ neurons were observed throughout the rostrocaudual extent of STN (Fig. 1*Q-S*). A high proportion (93.7 ± 0.8%) of these subthalamostriatal neurons (*n* = 314 total neurons counted from 4 mice) expressed nuclear immunoreactivity for FoxP2. Using unbiased stereological sampling methods, we estimated the total numbers of striatal starter neurons and Rb-GFP+ STN neurons in each VGAT-Cre:ChAT-Cre mouse (Fig. 1*I*). The average number of starter neurons per mouse was estimated to be 25,275 ± 2,820 whereas the average number of Rb-GFP+ STN neurons was 1,570 ± 137. To evaluate the relative connectivity from these stereological estimates, we calculated a ‘normalized connectivity index’ for each mouse (Do et al., 2016; Choi et al., 2019). On average, the normalized connectivity index in the VGAT-Cre:ChAT-Cre mice was 0.065 ± 0.010 (also see Fig. 6*A*). Thus, in general, striatal starter cells far outnumbered the retrogradely-labeled STN neurons. The results of these retrograde tracing studies suggest that glutamatergic STN neurons innervate and form synaptic connections with a host of neurons in the dorsal striatum.

### Subthalamic nucleus neurons rarely provide inputs to striatal spiny projection neurons

Striatal SPNs are heterogeneous in form and function. The direct/indirect pathways model of BG circuit organization (DeLong, 1990; Smith et al., 1998) posits a dichotomy in striatal output, instantiated by direct pathway SPNs (dSPNs) and indirect pathway SPNs (iSPNs). Monosynaptic retrograde tracing studies have revealed some quantitative differences in the extrinsic inputs to dorsal striatal dSPNs and iSPNs, although there are some clear inconsistencies (Wall et al., 2013; Guo et al., 2015; Fürth et al., 2018). To elucidate the extent to which STN neurons innervate dSPNs, we unilaterally and sequentially injected Cre-dependent helper virus and modified rabies virus into the dorsal striatum of adult Drd1a-Cre mice (*n* = 5; Fig. 2*A*). The majority of striatal starter neurons expressed Ctip2 (85.0 ± 5.6%; *n* = 2,261 total neurons counted from 4 mice). Of these Ctip2+ neurons, only a tiny fraction (0.6 ± 0.2%) also expressed somatic immunoreactivity for preproenkepahlin (PPE), a selective marker of iSPNs (Lee et al., 1997; Gerfen and Surmeier, 2011; Garas et al., 2016; Sharott et al., 2017). These data indicate a highly-selective transduction of dSPNs by the two viruses in the Drd1a-Cre mice. Focusing on STN, we then observed Rb-GFP+ neurons throughout its rostrocaudual extent (Fig. 2*B-D*). A high proportion (93.8 ± 1.6%) of these subthalamostriatal neurons (*n* = 223 total neurons counted from 5 mice) expressed FoxP2. To elucidate the extent to which STN neurons innervate iSPNs, we carried out the same monosynaptic retrograde tracing procedure in Adora2a-Cre mice (*n* = 5; Fig. 2*E*). The majority of striatal starter neurons expressed Ctip2 (94.8 ± 1.5%; *n* = 2,551 total neurons counted from 5 mice), and of these Ctip2+ neurons, almost all (97.7 ± 0.2%) co-expressed PPE, together indicating a highly-selective transduction of iSPNs in the Adora2a-Cre mice. Again, we observed Rb-GFP+ neurons throughout the rostrocaudual extent of STN (Fig. 2*F-H*), and a high proportion (95.4 ± 2.9%) of these subthalamostriatal neurons (*n* = 112 total neurons counted from 5 mice) expressed FoxP2. The average number of striatal starter neurons per Drd1a-Cre mouse was stereologically estimated to be 10,470 ± 2,597 whereas the average number of Rb-GFP+ STN neurons was 1,088 ± 350 (Fig. 2*I*). The average number of striatal starter neurons per Adora2a-Cre mouse was estimated to be 8,263 ± 2,139 whereas the average number of Rb-GFP+ STN neurons was 477 ± 124 (Fig. 2*I*). On average, the normalized connectivity index in Drd1a-Cre mice (0.098 ± 0.011) was significantly higher (*p* = 0.0183, unpaired t-test) than that in Adora2a-Cre mice (0.060 ± 0.007). Taken together, these anatomical data suggest that STN neurons modestly innervate both dSPNs and iSPNs, and that the innervation of dSPNs is comparatively greater.

**Figure 2.**
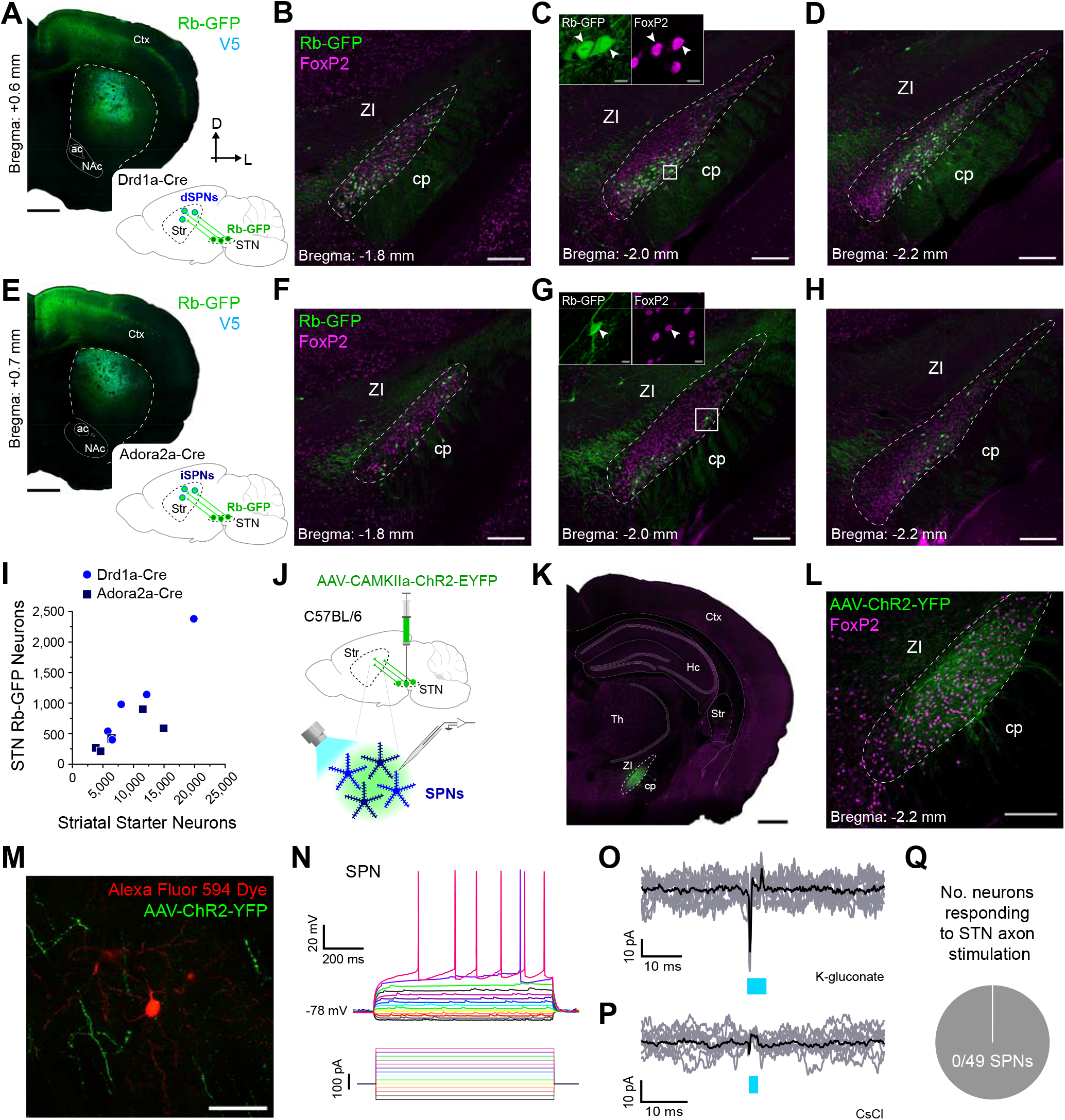
Subthalamic nucleus neurons rarely provide inputs to striatal spiny projection neurons. ***A***, Retrograde labeling of STN neurons that monosynaptically innervate direct pathway spiny projection neurons (dSPNs). *Inset*, a helper virus and modified rabies virus were unilaterally injected into the dorsal striatum of adult Drd1a-Cre mice. Neurons in STN that innervate starter dSPNs express rabies-encoded enhanced GFP (Rb-GFP, green). *Main image,* Coronal section from a Drd1a-Cre mouse, showing immunofluorescence signals for V5 (cyan) and Rb-GFP. Note the enriched co-localization of V5 and Rb-GFP signals, indicative of starter neurons, in the dorsal striatum (boundaries marked by dashed lines). ***B-D***, Retrogradely-labeled (Rb-GFP+) input neurons were located in rostral (***B***), central (***C***), and caudal (***D***) parts of the STN; all images from a single Drd1a-Cre mouse. Immunoreactivity for FoxP2 (magenta) was used to facilitate delineation of STN (dashed lines). ***C***, *Inset*, higher-magnification confocal image of Rb-GFP+ neurons within the white boxed area in ***C***; the Rb-GFP+ neurons often co-expressed FoxP2 (arrowheads). ***E***, Retrograde labeling of STN neurons that monosynaptically innervate indirect pathway SPNs (iSPNs). As for scheme in ***A***, but with use of Adora2a-Cre mice to target iSPNs. ***F-H***, Retrogradely-labeled input neurons were located in rostral (***F***), central (***G***), and caudal (***H***) parts of the STN; all images from a single Adora2a-Cre mouse. ***I***, Stereologically-estimated numbers of striatal starter neurons in Drd1a-Cre mice (*n* = 5) and Adora2a-Cre (*n* = 5) mice, respectively, as well as retrogradely-labeled STN neurons in these mice (each circle or square indicates estimates from a single Drd1a-Cre mouse or Adora2a-Cre mouse, respectively). ***J***, Main strategy for combined *ex vivo* electrophysiological and optogenetic interrogation of synaptic connections between STN neuron axons and striatal SPNs. A viral vector (AAV-CamKIIa-ChR2-EYFP) expressing the light-activated ion channel channelrhodopsin2 (ChR2) fused to a fluorescent reporter (EYFP) was first injected into the STN of wildtype (C57BL/6) mice. Visualized whole-cell patch-clamp recordings were then made from SPNs in acutely-prepared tissue slices from these mice, with brief pluses of blue light (470 nm) being used to selectively stimulate the axons of transduced ChR2-expressing STN neurons. ***K***, Coronal tissue section from a wildtype mouse, showing a circumscribed group of transduced neurons (AAV-ChR2-EYFP, green) in the STN (dashed lines). Hc; hippocampus; Th, thalamus. ***L***, Higher-magnification image of STN and neighboring brain regions from section shown in ***K. M***, SPN filled with Alexa Fluor 594 dye (red) during *ex vivo* whole-cell patch-clamp recordings, surrounded by ChR2-expressing axons of STN neurons (AAV-ChR2-EYFP, green). ***N***, Current-clamp recordings (K-gluconate-based pipette solution) of a neuron classified as an SPN, as per its characteristic voltage responses (*top*) to somatic injection of 900 ms pulses of hyperpolarizing or depolarizing current (*bottom*, from −100 pA to +225 pA, in 25 pA steps). ***O***, Voltage-clamp recordings (V_h_ = −70 mV) of an SPN, patched with K-gluconate solution, that showed no responses to optical stimulation of ChR2-expressing STN axons; nine individual trial traces (grey) are overlaid with an average trace (black). Blue bar indicates light on (5 ms flashes). The brief (<0.5 ms) deflections occurring at the onset and offset of light flashes are artifacts. ***P***, Voltage-clamp recordings (V_h_ = −70 mV) of another SPN, patched with CsCl-based pipette solution, showing no responses to the blue light flashes (2 ms). ***Q***, Pie chart showing the proportion of recorded SPNs that responded to optical stimulation of STN axons in the striatum; in this case, none of the 49 SPNs responded. Scale bars: ***A***, 750 μm; ***B-D***, 200 μm; ***C*** (inset), 10 μm; ***E***, 750 μm; ***F-H***, 750 μm; ***G*** (inset), 10 μm; ***K***, 750 μm; ***L***, 100 μm; ***M***, 50 μm.

To gain further insight into the properties of STN inputs to SPNs, we utilized *ex vivo* electrophysiology and optogenetics to test for neuronal responses to selective activation of subthalamostriatal axons. Specifically, we made visualized whole-cell patch-clamp recordings from SPNs in dorsal striatal slices from adult wildtype mice that had received STN injections of an AAV expressing the light-activated ion channel channelrhodopsin2 (ChR2) fused to enhanced yellow fluorescent protein (EYFP) under the control of the *Camk2a* (CaMKIIa) promoter (Fig. 2*J*). We then delivered single, brief flashes (2 or 5 ms) of blue light (470 nm, ~1.5 mW/mm^2^) to the slices to evoke neurotransmitter release from ChR2-expressing STN axons (Fig. 2*J*). This optogenetics-based strategy allowed for well-circumscribed transduction of STN neurons with ChR2, as revealed with EYFP (Fig. 2*K,L*). The identities of recorded SPNs were verified according to morphological criteria (as revealed after cell filling with fluorescent dye or biocytin; Fig. 2*M*) and, in cases where we used a K-gluconate-based pipette solution for recordings (see below), additionally confirmed by their distinctive intrinsic membrane properties (Fig. 2*N*; (Gertler et al., 2008)). We took care to ensure that each neuron was recorded in an area of the striatal slice that was innervated by ChR2-expressing axons (Fig. 2*M*). In this first level of analysis, we did not attempt to further classify the recorded neurons as dSPNs or iSPNs. We recorded 29 SPNs (from 6 mice) in both current-clamp mode and voltage-clamp mode (V_h_ −70 mV) using a K-gluconate pipette solution. Upon delivery of blue light flashes to the surrounding striatal tissue, none of these SPNs exhibited evoked responses (EPSPs or EPSCs, respectively) at latencies of <12 ms that should capture monosynaptic and polysynaptic responses (Fig. 2*O*). In using a K-gluconate pipette solution, we were mindful of the possibility that inadequate ‘space clamping’ could lead to false negatives, such that excitatory inputs to the distal dendrites of neurons might not be readily detected with recordings at their somata (Koós et al., 2004). To help address this, we recorded another 20 SPNs (from 4 mice) in voltage-clamp mode using a CsCl-based pipette solution that reduces electronic distance effects (Koós et al., 2004). Again, none of these SPNs exhibited monosynaptic or polysynaptic responses to the blue light flashes (Fig. 2*F*). In summary, 0 of 49 recorded SPNs (from 10 mice) exhibited a detectable response to selective activation of subthalamostriatal axons (Fig. 2*Q*; also see Fig. 6*B*). These electrophysiological data indicate that STN inputs to striatal spiny projection neurons are rare and, when assessed by somatic recordings, poorly efficacious.

### Subthalamic nucleus neurons commonly provide robust and reliable inputs to striatal parvalbumin-expressing interneurons

Monosynaptic retrograde tracing has revealed some quantitative differences in the glutamatergic cortical and thalamic inputs to dorsal striatal dSPNs and PV interneurons (Choi et al., 2019), raising the possibility that STN inputs to these two striatal cell types might also differ. To elucidate the extent to which STN neurons innervate PV interneurons, we unilaterally and sequentially injected Cre-dependent helper virus and modified rabies virus into the dorsal striatum of adult PV-Cre mice (*n* = 9; Fig. 3*A*). The majority of striatal starter neurons expressed somatic immunoreactivity for PV (95.5 ± 1.3%; *n* = 357 total neurons counted from 9 mice), attesting to a highly-selective transduction of PV interneurons by the two viruses. We then observed Rb-GFP+ neurons throughout the rostrocaudual extent of STN (Fig. 3*B-D*). A high proportion (90.7 ± 3.5%) of these subthalamostriatal neurons (*n* = 112 total neurons counted from 9 mice) expressed FoxP2. The average number of striatal starter neurons per PV-Cre mouse was stereologically estimated to be 640 ± 107 whereas the average number of Rb-GFP+ STN neurons was 262 ± 60 (Fig. 3*E*). On average, the normalized connectivity index in PV-Cre mice was relatively high (0.430 ± 0.067; also see Fig. 6*A*). Taken together, these anatomical data suggest that STN neurons robustly innervate striatal PV interneurons.

**Figure 3.**
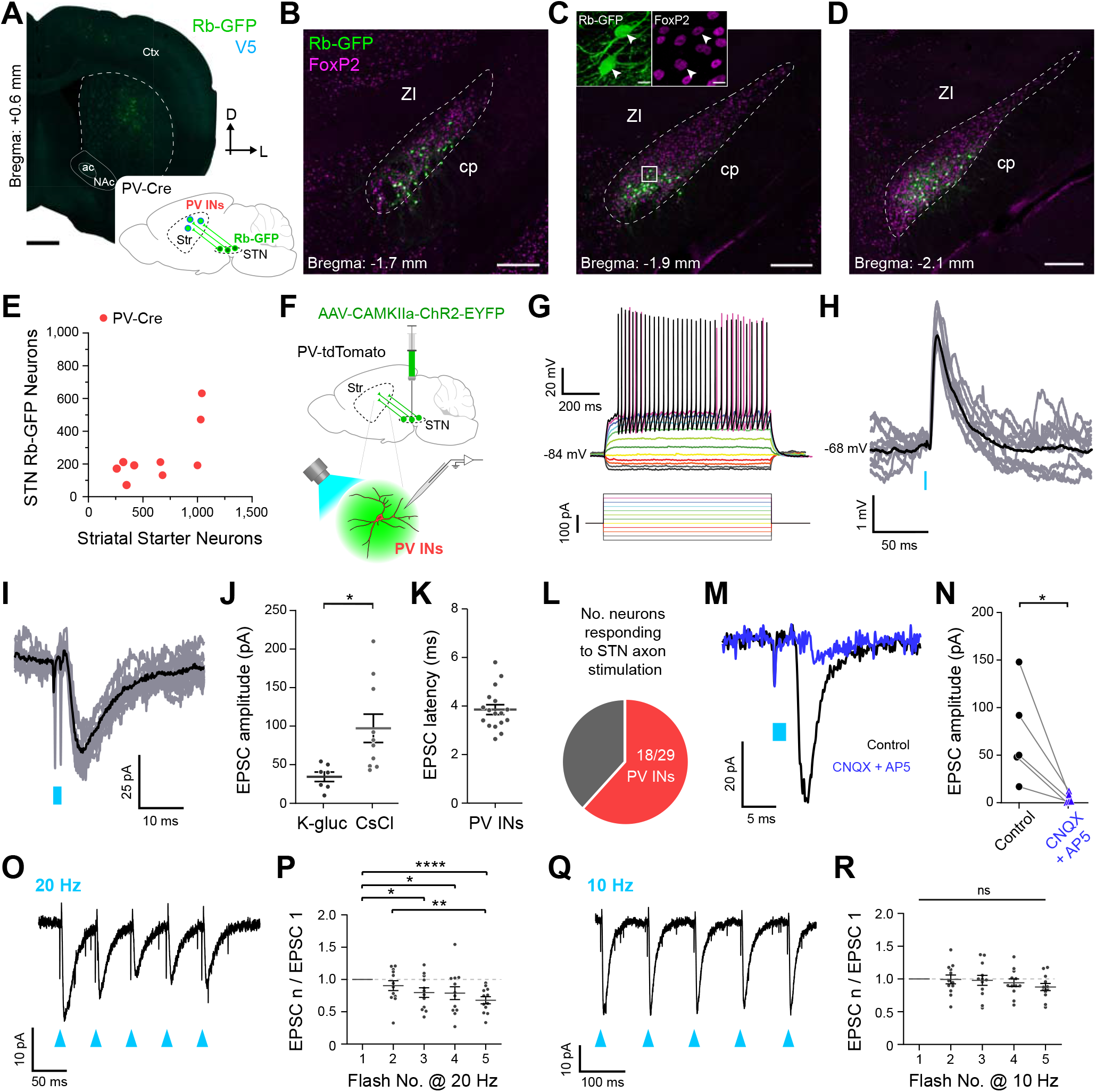
Subthalamic nucleus neurons provide robust and reliable inputs to striatal parvalbumin-expressing interneurons. ***A***, Retrograde labeling of STN neurons that monosynaptically innervate parvalbumin-expressing interneurons (PV INs). *Inset,* a helper virus and modified rabies virus were unilaterally injected into the dorsal striatum of adult PV-Cre mice. Neurons in STN that innervate starter PV INs express rabies-encoded enhanced GFP (Rb-GFP, green). *Main image,* Coronal section from a PV-Cre mouse, showing immunofluorescence signals for V5 (blue) and Rb-GFP. ***B-D***, Retrogradely-labeled (Rb-GFP+) input neurons in rostral (***B***), central (***C***), and caudal (***D***) parts of the STN; all images from a single PV-Cre mouse. ***C***, *Inset,* higher-magnification confocal image of Rb-GFP+ neurons within the white boxed area in ***C***; the Rb-GFP+ neurons often co-expressed FoxP2 (arrowheads). ***E***, Stereologically-estimated numbers of striatal starter neurons and retrogradely-labeled STN neurons in PV-Cre mice (*n* = 9). ***F***, Strategy for combined *ex vivo* electrophysiological and optogenetic interrogation of synaptic connections between STN neuron axons and striatal PV INs. A viral vector expressing channelrhodopsin2 (AAV-CamKIIa-ChR2-EYFP) was first injected into the STN of PV-tdTomato mice. Visualized whole-cell patch-clamp recordings were then made from tdTomato-expressing PV INs in tissue slices from these mice, with brief flashes of blue light (470 nm) being used to selectively stimulate the axons of transduced ChR2-expressing STN neurons. ***G***, Current-clamp recordings (K-gluconate-based pipette solution) of a PV IN, showing its characteristic voltage responses (*top*) to somatic injection of 900 ms pulses of hyperpolarizing or depolarizing current *(bottom,* from −100 pA to +175 pA, in 25 pA steps). ***H***, Current-clamp recordings of a PV IN, patched with K-gluconate pipette solution, showing consistent excitatory postsynaptic potentials in response to optical stimulation of ChR2-expressing STN axons; nine individual trial traces (grey) are overlaid with an average trace (black), with an indicative membrane potential. Blue bar indicates light on (2 ms flashes). ***I***, Voltage-clamp recordings (V_h_ = −70 mV) of the same PV IN as shown in ***H***, showing consistent excitatory postsynaptic currents (EPSCs) in response to blue light flashes (2 ms). ***J***, Peak amplitudes of evoked EPSCs for PV INs recorded with K-gluconate-based and CsCl-based pipette solutions (*n* = 7 and *n* =10 PV INs, respectively; unpaired t-test, \**p* = 0.015). ***K***, Latencies to onsets of EPSCs evoked in PV INs by STN axon stimulation (*n* = 17 PV INs, all at V_h_ = −70 mV). ***L***, Pie chart showing the proportion of recorded PV INs that responded to optical stimulation of STN axons in the striatum; in this case, 18 out of 29 PV INs responded. ***M***, Voltage-clamp recordings of another PV IN, patched with CsCl-based pipette solution, showing average responses to STN axon stimulation (from 10 individual trials) in control conditions (black trace), and after bath application of a combination of the AMPA/kainate-type glutamate receptor antagonist CNQX (10 μM) and the NMDA-type glutamate receptor antagonist AP5 (10 μM) (dark blue trace). ***N***, Pharmacological blockade of ionotropic glutamatergic receptors significantly attenuated the ESPCs evoked in PV INs (*n* = 5) by STN axon stimulation (paired t-test, \**p* = 0.0341). Neurons were recorded with either CsCl-based (*n* = 3) or K-gluconate-based pipette solution (*n* = 2). ***O***, EPSCs recorded from a PV IN (V_h_ = −70 mV; same neuron as ***H, I***) in response to a 20 Hz train of five light flashes (each of 2 ms; blue arrowheads). ***P***, Ratio of EPSCs evoked by flashes 2, 3, 4 or 5 to the EPSC evoked by the first flash during 20 Hz trains of STN axon stimulation (*n* = 12 PV INs). Over the course of the 5 flashes, there was on average a significant depression of the evoked responses (one-way repeated measures ANOVA [F(4,44) = 7.752, *p* = 0.000081] with Tukey’s *post hoc* tests [\**p* < 0.05; \*\**p* < 0.01; \*\*\*\**p* < 0.0001]). ***Q***, EPSCs recorded from a PV IN (V_h_ = −70 mV; same neuron as ***H, I, O***) in response to a 10 Hz train of five light flashes (each of 2 ms). ***R***, Ratio of EPSCs evoked by flashes 2, 3, 4 or 5 to the EPSC evoked by the first flash during 10 Hz trains of STN axon stimulation (*n* = 12 PV INs). Over the course of the 5 flashes, there were no significant changes in the evoked responses (one-way repeated measures ANOVA [F(4,44) = 1.621, *p* = 0.186]). Data in ***J, K, P*** and ***R*** are means ± SEMs, with each dot indicating data from a single neuron. Scale bars: ***A***, 750 μm; ***B-D***, 200 μm; ***C*** (inset), 10 μm.

To gain further insight into the properties of STN inputs to PV interneurons, we made whole-cell patch-clamp recordings from PV interneurons, as visualized by their native red fluorescence in striatal slices from adult mice expressing the reporter protein tdTomato under the control of the *Pvalb* promoter (PV-tdTomato mice), and tested their responses to optical activation of subthalamostriatal axons expressing ChR2 fused with EYFP (Fig. 3*F*). When recorded in current-clamp mode using a K-gluconate pipette solution, PV interneurons (*n* = 14, from 5 mice) exhibited intrinsic membrane properties typical of this cell type, *i.e.* conforming to a ‘fast spiking’ phenotype (Fig. 3*G*; (Tepper et al., 2010)). Upon delivery of single brief flashes (2 or 5 ms) of blue light, 8 of 14 PV interneurons recorded using K-gluconate consistently exhibited short-latency EPSPs (Fig. 3*H*) and/or EPSCs (Fig. 3*I*). The average peak EPSP amplitude was 1.55 ± 0.42 mV (*n* = 6 responsive PV interneurons), and did not cause the interneurons to fire action potentials. The average peak EPSC amplitude was 35.22 ± 7.31 pA (*n* = 7 responsive PV interneurons with V_h_ = −70 mV; Fig. 1*J*). We recorded another 15 PV interneurons (from 3 mice) in voltage-clamp mode using a CsCl-based pipette solution. The blue light flashes evoked robust EPSCs in 10 of 15 PV interneurons; the average peak EPSC amplitude was 97.14 ± 18.40 pA (Fig. 1*J*). The average latency to the onset of evoked EPSCs was 3.90 ± 0.21 ms (*n* = 17 responsive PV interneurons; Fig. 1*K*); because of the short-latency onsets of the evoked EPSCs, we considered all these responses of PV interneurons to be putatively monosynaptic (see Methods). In summary, 18 of 29 recorded PV interneurons (from 5 mice) exhibited clear and consistent responses to the selective activation of subthalamostriatal axons (Fig. 3*L*; also see Fig. 6*B*). The glutamatergic nature of the inputs from STN was verified in a subset of voltage-clamp recordings (*n* = 5 PV interneurons tested); bath application of antagonists of AMPA/kainate-type and NMDA-type ionotropic glutamate receptors (CNQX and AP5, respectively, both at 10 μM) greatly diminished the amplitudes of evoked EPSCs (Fig. 3*M,N*). Finally, we explored the short-term plasticity profiles of these subthalamostriatal inputs by delivering trains of 5 light flashes at inter-flash intervals of 50 ms or 100 ms (12 responsive PV interneurons; Fig. 3*O-R*). As compared to the EPSCs evoked by the first flashes in the trains, the amplitudes of subsequent EPSCs were successively reduced with optical stimulation at 20 Hz (Fig. 3*O,P*) but not at 10 Hz (Fig. 3*Q,R*). Taken together, these electrophysiological data indicate that STN inputs to striatal PV-expressing interneurons are relatively common and efficacious.

### Subthalamic nucleus neurons rarely provide inputs to two types of striatal neuropeptide Y-expressing interneurons

In rodent striatum, GABAergic interneurons expressing neuropeptide Y (NPY) are comprised of at least two major cell types that can be distinguished by a host of properties (Tepper et al., 2010, 2018). The most common type of NPY interneuron is defined by, amongst other features, its co-expression of somatostatin and nitric oxide synthase. Because these NPY/SOM/NOS interneurons exhibit characteristic low-threshold spikes (LTS) *ex vivo*, they are also commonly termed ‘LTS’ interneurons (Tepper et al., 2010, 2018). Dorsal striatal SOM and PV interneurons exhibit dissimilarities in their glutamatergic cortical and thalamic inputs (Assous and Tepper, 2019; Choi et al., 2019), again raising the possibility that STN inputs to these two striatal cell types might also differ. To elucidate the extent to which STN neurons innervate SOM interneurons, we unilaterally and sequentially injected Cre-dependent helper virus and modified rabies virus into the dorsal striatum of adult SOM-Cre mice (*n* = 5; Fig. 4*A*). The majority of striatal starter neurons expressed somatic immunoreactivity for somatostatin (98.0 ± 1.0%; *n* = 481 total neurons counted from 5 mice), attesting to a highly-selective transduction of SOM interneurons by the two viruses. Of these SOM-expressing interneurons, a large fraction (96.7 ± 1.7%) also expressed immunoreactivity for NOS, indicating that the vast majority of starter neurons likely correspond to LTS interneurons. We then observed Rb-GFP+ neurons throughout the rostrocaudual extent of STN (Fig. 4*B-D*). A high proportion (97.8 ± 2.2%) of these subthalamostriatal neurons (*n* = 49 total neurons counted from 5 mice) expressed FoxP2. The average number of striatal starter neurons per SOM-Cre mouse was stereologically estimated to be 2,019 ± 216 whereas the average number of Rb-GFP+ STN neurons was 196 ± 47 (Fig. 4*F*). On average, the normalized connectivity index in SOM-Cre mice was 0.098 ± 0.025 (also see Fig. 6*A*). Taken together, these anatomical data suggest that STN neurons modestly innervate striatal NPY/SOM/NOS-expressing LTS interneurons.

**Figure 4.**
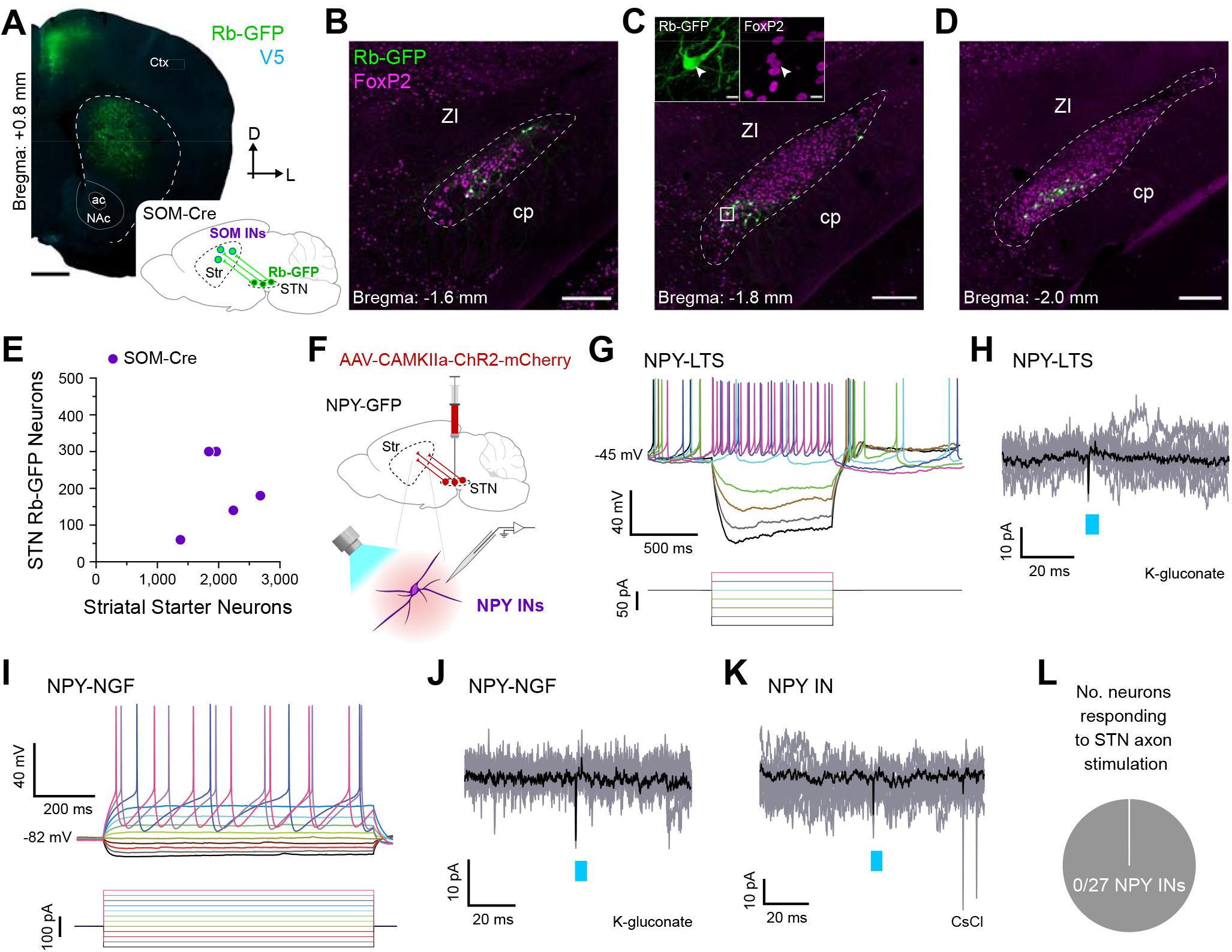
Subthalamic nucleus neurons rarely provide inputs to two types of striatal neuropeptide Y-expressing striatal interneurons. ***A***, Retrograde labeling of STN neurons that monosynaptically innervate somatostatin-expressing interneurons (SOM INs), many of which also express neuropeptide Y (NPY). *Inset,* a helper virus and modified rabies virus were unilaterally injected into the dorsal striatum of adult SOM-Cre mice. Neurons in STN that innervate starter SOM INs express rabies-encoded enhanced GFP (Rb-GFP, green). *Main image,* Coronal section from a SOM-Cre mouse, showing immunofluorescence signals for V5 (blue) and Rb-GFP. ***B-D***, Retrogradely-labeled (Rb-GFP+) input neurons in rostral (***B***), central (***C***), and caudal (***D***) parts of the STN; all images from a single SOM-Cre mouse. ***C***, *Inset,* higher-magnification confocal image of Rb-GFP+ neurons within the white boxed area in ***C***; the Rb-GFP+ neurons often co-expressed FoxP2 (arrowheads). ***E***, Stereologically-estimated numbers of striatal starter neurons and retrogradely-labeled STN neurons in SOM-Cre mice (*n* = 5). ***F***, Main strategy for combined *ex vivo* electrophysiological and optogenetic interrogation of synaptic connections between STN neuron axons and striatal NPY-expressing interneurons (NPY INs). A viral vector expressing channelrhodopsin2 (AAV-CamKIIa-ChR2-mCherry) was first injected into the STN of NPY-GFP mice. Visualized whole-cell patch-clamp recordings were then made from GFP-expressing (NPY) INs in tissue slices from these mice, with brief flashes of blue light (470 nm) being used to selectively stimulate the axons of transduced ChR2-expressing STN neurons. ***G***, Current-clamp recordings (K-gluconate-based pipette solution) of a NPY IN, showing its characteristic voltage responses (*top*) to somatic injection of 900 ms pulses of hyperpolarizing or depolarizing current (*bottom,* from −100 pA to +50 pA, in 25 pA steps). Note this cell exhibited low-threshold spike bursts soon after the cessation of some hyperpolarizing current pulses and thus, was classified as a NPY-LTS interneuron. ***H***, Voltage-clamp recordings (V_h_ = −70 mV) of a NPY-LTS interneuron (same neuron as shown in ***G***), showing no responses to optical stimulation of ChR2-expressing STN axons; nine individual trial traces (grey) are overlaid with an average trace (black). Blue bar indicates light on (5 ms flashes). ***I***, Current-clamp recordings (K-gluconate-based pipette solution) of a NPY IN classified as a neurogliaform (NGF) interneuron, as per its characteristic voltage responses (*top*) to somatic injection of 900 ms pulses of hyperpolarizing or depolarizing current (*bottom*, from −100 pA to +175 pA, in 25 pA steps). ***J***, Voltage-clamp recordings (V_h_ = −70 mV) of a NPY-NGF interneuron (same neuron as shown in ***I***), showing no responses to optical stimulation of ChR2-expressing STN axons. ***K***, Voltage-clamp recordings (V_h_ = −70 mV) of another NPY IN, patched with CsCl-based pipette solution, showing no responses to the blue light flashes (5 ms). ***L***, Pie chart showing the proportion of recorded NPY INs that responded to optical stimulation of STN axons in the striatum; in this case, none of the 27 NPY INs responded (samples were 14 LTS INs and 4 NGF INs recorded using K-gluconate based pipette solution, and 9 other INs recorded using CsCl-based pipette solution). Scale bars: ***A***, 750 μm; ***B-D***, 200 μm; ***C*** (inset), 10 μm.

To gain further insight into these neuronal connections, we made whole-cell patchclamp recordings from LTS interneurons, as visualized by their native green fluorescence in striatal slices from adult mice expressing humanized *Renilla* GFP under the control of the *Npy* promoter (NPY-GFP mice; (Ibáñez-Sandoval et al., 2011)), and tested their responses to optical activation of subthalamostriatal axons expressing ChR2 fused with the reporter protein mCherry (Fig. 4*F*). When recorded in current-clamp mode using a K-gluconate pipette solution, NPY-expressing LTS interneurons (*n* = 14, from 3 mice) were readily classified as such by their characteristic membrane properties (Fig. 4*G*; (Ibáñez-Sandoval et al., 2011)). When recorded in current-clamp or voltage-clamp modes, none of these LTS interneurons exhibited monosynaptic or polysynaptic responses to the local delivery of brief flashes of blue light (Fig. 4*H*).

The second major type of NPY-expressing cell in mouse striatum, the so-called neurogliaform (NGF) interneuron, does not express SOM or NOS, and does not exhibit LTS activity (Ibáñez-Sandoval et al., 2011). Our use of SOM-Cre mice for monosynaptic retrograde tracing allowed for highly-selective access to LTS interneurons but, in doing so, precluded structural analyses of STN inputs to NGF interneurons. Previous work has shown that LTS interneurons and NGF interneurons differ considerably in their responses to cortical and thalamic inputs (Assous et al., 2017; Assous and Tepper, 2019). It is not known whether the responses of LTS and NGF interneurons to STN inputs also differ. To address this, we also made whole-cell patch-clamp recordings from NGF interneurons, as visualized in striatal slices from adult NPY-GFP mice, and tested their responses to optical activation of ChR2-expressing subthalamostriatal axons (Fig. 4*F*). When recorded in current-clamp mode using a K-gluconate pipette solution, NPY-expressing NGF interneurons (*n* = 4, from 2 mice) were readily classified as such by their distinctive membrane properties (Fig. 4*I*; (Ibáñez-Sandoval et al., 2011)). When recorded in current-clamp or voltage-clamp mode, none of these NGF interneurons responded to the delivery of brief flashes of blue light (Fig. 4*J*).

In a final set of experiments, we recorded another 9 NPY interneurons (from 3 mice) in voltage-clamp mode using a CsCl-based pipette solution (note this solution precluded their electrophysiological classification as LTS or NGF interneurons). Again, none of these interneurons responded to the blue light flashes (Fig. 4*K*). In summary, 0 of 27 recorded NPY interneurons (from 4 mice) exhibited a detectable response to selective activation of subthalamostriatal axons (Fig. 4*L*; also see Fig. 6*B*). These electrophysiological data indicate that STN inputs to striatal neuropeptide Y-expressing interneurons are rare and, when assessed by somatic recordings, poorly efficacious.

### Subthalamic nucleus neurons rarely provide inputs to striatal cholinergic interneurons

Monosynaptic retrograde tracing has revealed some quantitative differences in the glutamatergic cortical and thalamic inputs to dorsal striatal cholinergic interneurons as compared to SPNs (Guo et al., 2015) and PV interneurons (Klug et al., 2018), highlighting the prospect that STN inputs to cholinergic interneurons might also differ. To elucidate the extent to which STN neurons innervate cholinergic interneurons, we unilaterally and sequentially injected Cre-dependent helper virus and modified rabies virus into the dorsal striatum of adult ChAT-Cre mice (*n* = 5; Fig. 5*A*). The majority of striatal starter neurons expressed somatic immunoreactivity for ChAT (90.9 ± 3.6%; *n* = 513 total neurons counted from 5 mice), confirming a highly-selective transduction of cholinergic interneurons by the two viruses. We then observed sparse Rb-GFP+ neurons scattered throughout the rostrocaudual extent of STN (Fig. 5*B-D*). A high proportion (82.1 ± 6.0%) of these subthalamostriatal neurons (*n* = 26 total neurons counted from 5 mice) expressed FoxP2.

**Figure 5.**
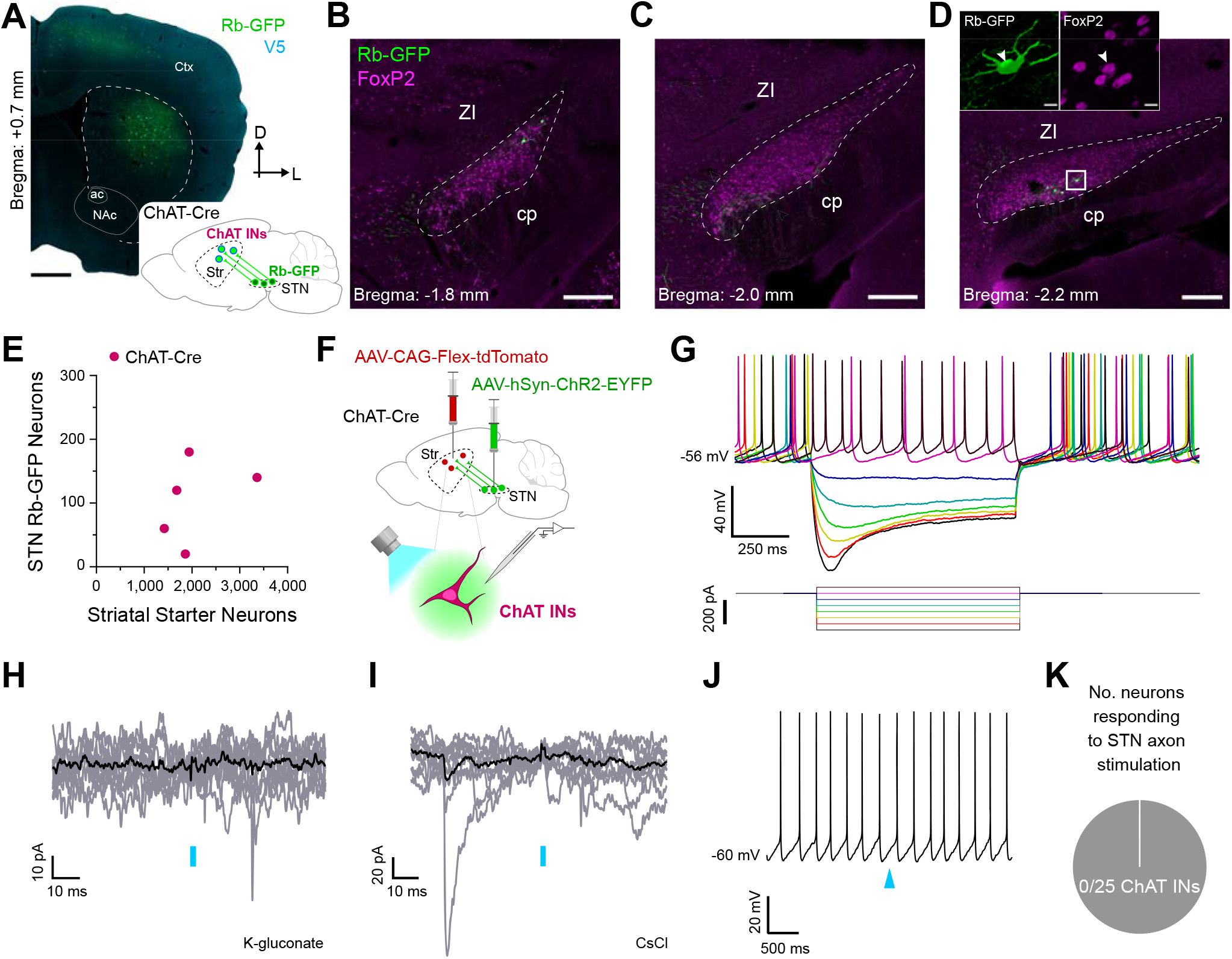
Subthalamic nucleus neurons rarely provide inputs to striatal cholinergic interneurons. ***A***, Retrograde labeling of STN neurons that monosynaptically innervate cholinergic interneurons (ChAT INs). *Inset,* a helper virus and modified rabies virus were unilaterally injected into the dorsal striatum of adult ChAT-Cre mice. Neurons in STN that innervate starter ChAT INs express rabies-encoded enhanced GFP (Rb-GFP, green). *Main image,* Coronal section from a ChAT-Cre mouse, showing immunofluorescence signals for V5 (blue) and Rb-GFP. ***B-D***, Retrogradely-labeled (Rb-GFP+) input neurons in rostral (***B***), central (***C***), and caudal (***D***) parts of the STN; all images from a single ChAT-Cre mouse. ***D***, */nset,* higher-magnification confocal image of Rb-GFP+ neurons within the white boxed area in ***D***; the Rb-GFP+ neurons often co-expressed FoxP2 (arrowheads). ***E***, Stereologically-estimated numbers of striatal starter neurons and retrogradely-labeled STN neurons in ChAT-Cre mice (*n* = 5). ***F***, Main strategy for combined *ex vivo* electrophysiological and optogenetic interrogation of synaptic connections between STN neuron axons and striatal ChAT INs. A Cre-dependent viral vector expressing tdTomato (AAV-CAG-Flex-tdTomato) was injected into the dorsal striatum of ChAT-Cre mice. Another viral vector expressing channelrhodopsin2 (AAV-CamKIIa-ChR2-EYFP) was injected into the STN of the same animals. Visualized whole-cell patch-clamp recordings were then made from tdTomato-expressing (ChAT) INs in tissue slices from these mice, with brief flashes of blue light (470 nm) being used to selectively stimulate the axons of transduced ChR2-expressing STN neurons. ***G***, Current-clamp recordings (K-gluconate-based pipette solution) of a ChAT IN, showing its characteristic voltage responses (*top*) to somatic injection of 900 ms pulses of hyperpolarizing or depolarizing current (*bottom*, from −300 pA to +50 pA, in 50 pA steps). ***H***, Voltage-clamp recordings (V_h_ = −70 mV) of a ChAT IN, patched with a K-gluconate pipette solution, showing no responses to optical stimulation of ChR2-expressing STN axons; nine individual trial traces (grey) are overlaid with an average trace (black). Blue bar indicates light on (2 ms flashes). ***I***, Voltage-clamp recordings (V_h_ = −70 mV) of another ChAT IN, patched with CsCl-based pipette solution, showing no responses to the blue light flashes (2 ms). ***J***, Current-clamp recording of a spontaneously-active ChAT IN, showing no response to optical stimulation of STN axons. Blue arrowhead indicates light on (2 ms flash). ***K***, Pie chart showing the proportion of recorded ChAT INs that responded to optical stimulation of STN axons in the striatum; in this case, none of the 25 ChAT INs responded. Scale bars: ***A***, 750 μm; ***B-D***, 200 μm; ***D*** (inset), 10 μm.

The average number of striatal starter neurons per ChAT-Cre mouse was stereologically estimated to be 2,052 ± 339 whereas the average number of Rb-GFP+ STN neurons was 104 ± 29 (Fig. 5*E*). On average, the normalized connectivity index in ChAT-Cre mice was 0.052 ± 0.014 (also see Fig. 6*A*). Taken together, these anatomical data suggest that STN neurons modestly innervate striatal cholinergic interneurons.

**Figure 6.**
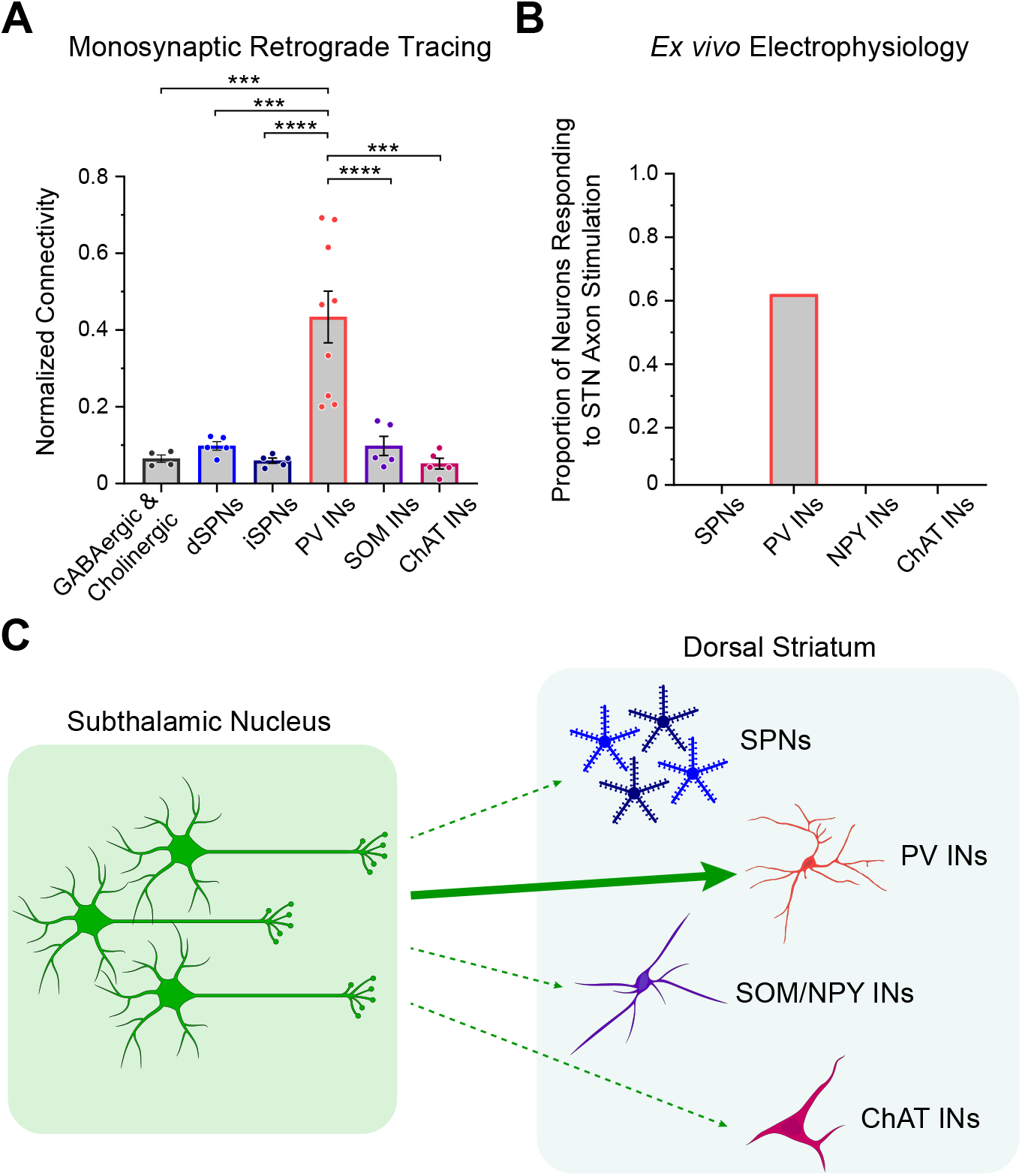
Subthalamic nucleus neurons selectively innervate striatal parvalbumin-expressing interneurons. ***A***, Summary of key data from monosynaptic retrograde tracing experiments. The normalized connectivity from the STN to striatal PV interneurons is significantly higher than that from STN to any of the four other striatal cell types examined (dSPNs, iSPNs, SOM INs, ChAT INs), as well as from STN to striatal neurons as a whole (GABAergic & Cholinergic; as studied using VGAT-Cre:ChAT-Cre mice). One-way ANOVA (F(5,27) = 13, *p* = 0.000001) with Tukey’s *post hoc* tests (\*\*\**p* < 0.001; **** *p* < 0.0001). Data are means ± SEMs, with each dot indicating data from a single mouse. ***B***, Summary of key results of *ex vivo* electrophysiological/optogenetics experiments. Of the five striatal cell types tested (SPNs, PV INs, NPY INs [includes both NPY-LTS, which are assumed to co-express SOM, and NPY-NGF types] and ChAT INs), only PV interneurons commonly exhibited responses to selective stimulation of subthalamostriatal axons. ***C***, Schematic representation of STN innervation of striatal cell types, taking into account both structural and electrophysiological readouts of connectivity. Thin dashed arrows indicate efferent connections supported by anatomical evidence but not by electrophysiological evidence. The thick solid arrow indicates strong, corroborative evidence of a monosynaptic connection. There is a highly selective and impactful glutamatergic projection from the STN to striatal parvalbumin-expressing interneurons.

To gain further insight, we made whole-cell patch-clamp recordings from cholinergic interneurons, as visualized by their native red fluorescence in striatal slices from adult ChAT-Cre mice that had received striatal injections of a Cre-dependent AAV expressing tdTomato, and tested their responses to optical activation of subthalamostriatal axons expressing ChR2 fused with EYFP (Fig. 5*F*). When recorded in current-clamp mode using a K-gluconate pipette solution, these cholinergic interneurons (*n* =17 neurons from 4 mice) exhibited intrinsic membrane properties typical of this cell type, including spontaneous firing in the absence of current injection (Fig. 5*G,J*; (Goldberg and Reynolds, 2011)). When recorded in current-clamp or voltage-clamp mode, none of these cholinergic interneurons exhibited monosynaptic or polysynaptic responses to the local delivery of brief flashes of blue light (Fig. 5*H,J*). We also recorded another 8 cholinergic interneurons (from 3 mice) in voltage-clamp mode using a CsCl-based pipette solution. Again, none of these interneurons responded to the optical stimuli (Fig. 5*I*). In summary, 0 of 25 recorded cholinergic interneurons (from 7 mice) exhibited a detectable response to selective activation of subthalamostriatal axons (Fig. 5*K*; also see Fig. 6*B*). These electrophysiological data indicate that STN inputs to striatal cholinergic interneurons are rare and, when assessed by somatic recordings, poorly efficacious.

### Comparative analyses of subthalamostriatal connections

A strength of our work here is the ability to directly compare the relative levels of STN innervation of multiple cell types in dorsal striatum, using both structural and electrophysiological readouts of connectivity. Taken together, the results of our monosynaptic retrograde tracing studies showed that the connectivity from STN neurons to striatal PV interneurons is significantly higher (~ four-to eight-fold) than that from STN to any of the four other striatal cell types examined, as well as from STN to striatal neurons as a whole (Fig. 6*A*). Of equal note, the connectivities of dSPNs, iSPNs, SOM interneurons and ChAT interneurons were quantitatively similar when compared altogether (Fig. 6*A*). The collective results of our *ex vivo* electrophysiological/optogenetics experiments showed that, of the five striatal cell types tested, only PV interneurons (and 62% of them) exhibited robust monosynaptic excitatory responses to selective activation of subthalamostriatal axons (Fig. 6*B*). We conclude that there is a highly selective and impactful glutamatergic projection from the STN to striatal PV interneurons (Fig. 6*C*).

## Discussion

Here, we provide structural and electrophysiological evidence of a remarkable target selectivity in the mouse subthalamostriatal projection. Our data converge to support the concept that glutamatergic STN neurons are positioned to directly and powerfully influence striatum by virtue of their enriched innervation of parvalbumin-expressing interneurons.

### Anatomical readouts of subthalamostriatal connectivity

Monosynaptic retrograde tracing from genetically-defined cell types has generated important new insights into the sources and organization of extrinsic inputs to striatum (Wall et al., 2013; Guo et al., 2015; Smith et al., 2016; Fürth et al., 2018; Klug et al., 2018; Monteiro et al., 2018; Choi et al., 2019; Melendez-Zaidi et al., 2019). Using this approach, we provide comparative analyses of the STN innervation of five cell types in dorsal striatum. We observed that the STN innervates dSPNs, iSPNs, PV interneurons, SOM (LTS) interneurons, and cholinergic interneurons. Importantly though, the connectivity from STN neurons to striatal PV interneurons stands apart in being of comparatively high magnitude. The precise substrates for this striking selectivity remain to be determined. A higher connectivity index could reflect one or more structural configurations, including (but not limited to): a relatively larger fraction of PV interneurons is innervated by STN; an individual PV interneuron is more likely to receive convergent inputs from multiple STN neurons; an individual STN neuron is more likely to form multiple synapses with a given PV interneuron, thus increasing the likelihood of presynaptic neuron transduction. Past studies of dorsal striatum employing monosynaptic retrograde tracing have compared ‘whole brain’ inputs to two or three striatal cell types, and some have reported that STN provides inputs to SPNs and/or interneurons (Wall et al., 2013; Guo et al., 2015; Smith et al., 2016; Klug et al., 2018; Choi et al., 2019). All these studies indicate that inputs from STN are relatively scant, at least when compared to other glutamatergic inputs from cortex and thalamus. None of these studies reported significant differences in the innervation of multiple striatal cell types by STN neurons. This apparent discordance with our results might arise from differences in tools used, the cells transduced for tracing, and/or the metrics analyzed. Unlike other studies, we: (1) injected a single helper virus, avoiding ambiguities in defining starter cells with all components necessary for retrograde labeling of their presynaptic partners; (2) quantified starter cell ‘specificity’ for all mouse lines; (3) employed unbiased stereology to estimate total numbers of starter cells and input neurons; and (4) used a normalized connectivity index focused on STN only, rather than expressing counts as a fraction of whole brain inputs. We conclude from our anatomical experiments that there is a highly-selective projection from the STN to striatal PV interneurons. Accordingly, the STN joins a growing list of subcortical structures that, although not considered canonical sources of inputs to striatum, selectively target striatal interneurons; exemplars include the GPe (Bevan et al., 1998; Mallet et al., 2012) and peduculopontine nucleus (Assous et al., 2019). We speculate that selective innervation of striatal interneurons is a common circuit motif.

### Electrophysiological readouts of subthalamostriatal connectivity

Building on the anatomical data above, we used a combination of *ex vivo* electrophysiology and optogenetics to gain the first direct insights into the incidence and strength of subthalamostriatal neurotransmission. We observed that, of the five striatal cell types tested, only PV interneurons commonly exhibited monosynaptic excitatory responses to activation of subthalamostriatal axons. The STN inputs to PV interneurons were glutamatergic, robust and reliable, with no or moderate attenuation when driven at frequencies similar to STN neuron firing rates *in vivo i.e.* 10-20 Hz (Mallet et al., 2008a, 2008b; Deffains et al., 2016). As such, the anatomical and electrophysiological readouts of connectivity were in broad agreement, in that both provided strong evidence of a highly-selective innervation of PV interneurons by the STN. However, the lack of responsive SPNs, NPY-LTS interneurons and cholinergic interneurons was at variance with anatomical data showing a modest innervation of these cell types. It is possible that STN inputs located at the distal dendrites of these striatal neurons were not detected by our recordings made at their somata, although our use of a CsCl-based pipette solution should have minimized electronic distance effects and chances of false negatives (Koós et al., 2004). Interestingly, there are other cases of mismatches between the anatomical connectivity, as defined by monosynaptic retrograde tracing, and electrophysiological connectivity of striatal circuits (Choi et al., 2019).

### Implications for striatal microcircuits

The efficacious monosynaptic connection from the STN to PV interneurons has important implications for the organization of activity in striatal microcircuits. GABAergic PV interneurons can exert powerful effects on their SPN targets, for example, delaying or negating action potential firing (Koós and Tepper, 1999; Tepper et al., 2004). It follows that suprathreshold excitation of PV interneurons by STN inputs could provide a novel substrate for ‘feed-forward’ inhibition in striatum. The efferent connections of PV interneurons in turn suggest that dSPNs, iSPNs and NPY/SOM/NOS-expressing LTS interneurons (and some PV interneurons) would be subject to this feed-forward inhibition (Gittis et al., 2010; Planert et al., 2010; Tepper et al., 2010, 2018; Szydlowski et al., 2013). Notably, cholinergic interneurons are not extensively targeted by PV interneurons (Szydlowski et al., 2013) or STN neurons.

### Subthalamostriatal combinatorics

The STN as a whole innervates all other BG nuclei, as well as discrete parts of midbrain, brainstem, thalamus and cerebral cortex (Smith et al., 1998; Emmi et al., 2020). Because STN neurons exhibit markedly heterogeneous structural (axonal), physiological and molecular properties (Bevan et al., 2000; Sato et al., 2000; Koshimizu et al., 2013; Wallén-Mackenzie et al., 2020), it seems unlikely that all STN neurons innervate all of these diverse targets. It is currently unclear whether all STN neurons innervating striatum also innervate, via axon collaterals, one or more other BG nuclei. Single-neuron tracings in rats suggest the majority of subthalamostriatal neurons additionally project to the GPe, entopeduncular nucleus and substantia nigra, but there are further combinations of projections (Koshimizu et al., 2013). Single-axon reconstructions in monkeys present another scenario in which about one fifth of STN neurons might innervate striatum but not other BG nuclei; we note however this interpretation is challenged by incomplete labeling of axon collaterals (Sato et al., 2000). Our anatomical data show that at least five types of striatal neuron receive inputs from STN, albeit to differing degrees, but it remains to be determined whether an individual subthalamostriatal neuron can innervate more than one type of striatal neuron; the potential number of combinations of subthalamostriatal connections is large. Our experiments were focused on central aspects of dorsal striatum (caudate-putamen) and, as such, preclude conclusions as to whether the same patterns of subthalamostriatal innervation also hold for all territories of dorsal striatum and/or any part of ventral striatum (nucleus accumbens). Notably, the functional properties of PV interneurons vary across striatal regions (Garas et al., 2016; Monteiro et al., 2018), and this heterogeneity might extend to their inputs from STN. Our anterograde tracing revealed a relatively sparse projection from STN to nucleus accumbens, in agreement with studies in rats (Groenewegen and Berendse, 1990; Koshimizu et al., 2013).

### Wider circuit context

The subthalamostriatal projection is not included in the direct/indirect pathways scheme nor (to our knowledge) any other model of the functional organization of cortical-basal ganglia-thalamocortical circuits. A feed-forward flow of information through these circuits is central to the direct/indirect pathways scheme (DeLong, 1990; Smith et al., 1998). Conceptually, the striatum lies ‘upstream’ of STN in the indirect pathway (Gerfen and Surmeier, 2011). One implication of this arrangement is that the routing back of STN output to striatum must occur through polysynaptic pathways, classically those incorporating the BG output nuclei and their thalamic effectors. There is scope for these sequential connections to degrade information carried by outgoing STN signals before it reaches striatum. In contrast, the monosynaptic projection from STN to striatum that we elucidate here offers a substrate by which striatal neurons, and particularly PV interneurons, can be quickly updated on STN activity dynamics with minimal distortion of signal. Given that STN selectively innervates GABAergic PV interneurons, which in turn innervate and powerfully control GABAergic iSPNs, this permutation of subthalamostriatal feedback could subserve a homeostatic function along the indirect pathway; increased STN firing would ultimately result in less iSPN output and thence, augmented GABAergic output from prototypic GPe neurons (Abdi et al., 2015) that would, in turn, restrain STN activity. The importance of appropriately constraining STN activity is exemplified in Parkinsonism where STN neurons are hyperactive (Mallet et al., 2008a, 2008b). Another permutation of subthalamostriatal feedback, mediated by STN innervation of dSPNs, presents a novel substrate for activity along the indirect pathway to influence that along the direct pathway. All variants of rapid monosynaptic subthalamostriatal feedback have implications for another influential model of BG functional organization that emphasizes the so-called hyperdirect pathway and is based on the cortical driving of STN neuron output before SPN output (Nambu et al., 2002). We conclude that the cell-type-selective innervation of striatum by glutamatergic STN neurons, as detailed here, is positioned to fulfil diverse and likely unique roles within BG circuits.

## Acknowledgements

The work of K.K. and T.K. was supported by National Institutes of Health grant NS034865. The work of N.M.D., O.A. and P.J.M. was supported by the UK Medical Research Council (Awards MC_UU_12024/2 and MC_UU_00003/5 to P.J.M.) and the Wellcome Trust (Investigator Award 101821 to P.J.M.). The work of K.M. was supported by the Swedish Research Council and the Swedish Brain Foundation (Hjärnfonden). We thank M. Assous, J. Tepper and D. Dautan for insightful scientific discussions, and J. Westcott, L. Conyers, H. Zhang, B. Micklem, F. Shah and O. Tzortzi for technical assistance.

## Author contributions

K.K., N.M.D., K.M., P.J.M. and T.K. designed research; K.K., N.M.D., O.A., B.K., A.T. and D.C. performed research; K.K., N.M.D. and O.A. analyzed data; K.K., N.M.D., P.J.M. and T.K. wrote the paper.

## Notes

### Competing Interest Statement

The authors have declared no competing interest.

## References

Abdi A, Mallet N, Mohamed FY, Sharott A, Dodson PD, Nakamura KC, Suri S, Avery SV, Larvin JT, Garas FN, Garas SN, Vinciati F, Morin S, Bezard E, Baufreton J, Magill PJ (2015) Prototypic and arkypallidal neurons in the dopamine-intact external globus pallidus. J Neurosci 35:6667–6688.

Ährlund-Richter S, Xuan Y, van Lunteren JA, Kim H, Ortiz C, Pollak Dorocic I, Meletis K, Carlén M (2019) A whole-brain atlas of monosynaptic input targeting four different cell types in the medial prefrontal cortex of the mouse. Nat Neurosci 22:657–668.

Albin RL, Aldridge JW, Young AB, Gilman S (1989) Feline subthalamic nucleus neurons contain glutamate-like but not GABA-like or glycine-like immunoreactivity. Brain Res 491:185–188.

Arlotta P, Molyneaux BJ, Jabaudon D, Yoshida Y, Macklis JD (2008) Ctip2 controls the differentiation of medium spiny neurons and the establishment of the cellular architecture of the striatum. The Journal of Neuroscience: The Official Journal of the Society for Neuroscience 28:622–632.

Assous M, Dautan D, Tepper JM, Mena-Segovia J (2019) Pedunculopontine Glutamatergic Neurons Provide a Novel Source of Feedforward Inhibition in the Striatum by Selectively Targeting Interneurons. J Neurosci 39:4727–4737.

Assous M, Kaminer J, Shah F, Garg A, Koós T, Tepper JM (2017) Differential processing of thalamic information via distinct striatal interneuron circuits. Nature Communications 8:15860.

Assous M, Tepper JM (2019) Excitatory extrinsic afferents to striatal interneurons and interactions with striatal microcircuitry. Eur J Neurosci 49:593–603.

Barroso-Chinea P, Castle M, Aymerich MS, Pérez-Manso M, Erro E, Tuñon T, Lanciego JL (2007) Expression of the mRNAs encoding for the vesicular glutamate transporters 1 and 2 in the rat thalamus. J Comp Neurol 501:703–715.

Beckstead RM (1983) A reciprocal axonal connection between the subthalamic nucleus and the neostriatum in the cat. Brain Res 275:137–142.

Bevan MD, Booth PA, Eaton SA, Bolam JP (1998) Selective innervation of neostriatal interneurons by a subclass of neuron in the globus pallidus of the rat. J Neurosci 18:9438–9452.

Bevan MD, Wilson CJ, Bolam JP, Magill PJ (2000) Equilibrium potential of GABA(A) current and implications for rebound burst firing in rat subthalamic neurons in vitro. Journal of Neurophysiology 83:3169–3172.

Callaway EM, Luo L (2015) Monosynaptic Circuit Tracing with Glycoprotein-Deleted Rabies Viruses. J Neurosci 35:8979–8985.

Choi K, Holly EN, Davatolhagh MF, Beier KT, Fuccillo MV (2019) Integrated anatomical and physiological mapping of striatal afferent projections. Eur J Neurosci 49:623–636.

Deffains M, Iskhakova L, Katabi S, Haber SN, Israel Z, Bergman H (2016) Subthalamic, not striatal, activity correlates with basal ganglia downstream activity in normal and parkinsonian monkeys. eLife 5.

DeLong MR (1990) Primate models of movement disorders of basal ganglia origin. Trends Neurosci 13:281–285.

Do JP, Xu M, Lee S-H, Chang W-C, Zhang S, Chung S, Yung TJ, Fan JL, Miyamichi K, Luo L, Dan Y (2016) Cell type-specific long-range connections of basal forebrain circuit. Elife 5.

Dodson PD, Larvin JT, Duffell JM, Garas FN, Doig NM, Kessaris N, Duguid IC, Bogacz R, Butt SJB, Magill PJ (2015) Distinct developmental origins manifest in the specialized encoding of movement by adult neurons of the external globus pallidus. Neuron 86:501–513.

Emmi A, Antonini A, Macchi V, Porzionato A, De Caro R (2020) Anatomy and Connectivity of the Subthalamic Nucleus in Humans and Non-human Primates. Frontiers in Neuroanatomy 14:13.

Faust TW, Assous M, Tepper JM, Koós T (2016) Neostriatal GABAergic Interneurons Mediate Cholinergic Inhibition of Spiny Projection Neurons. The Journal of Neuroscience: The Official Journal of the Society for Neuroscience 36:9505–9511.

Fürth D, Vaissière T, Tzortzi O, Xuan Y, Märtin A, Lazaridis I, Spigolon G, Fisone G, Tomer R, Deisseroth K, Carlén M, Miller CA, Rumbaugh G, Meletis K (2018) An interactive framework for whole-brain maps at cellular resolution. Nat Neurosci 21:139–149.

Garas FN, Shah RS, Kormann E, Doig NM, Vinciati F, Nakamura KC, Dorst MC, Smith Y, Magill PJ, Sharott A (2016) Secretagogin expression delineates functionally-specialized populations of striatal parvalbumin-containing interneurons. Elife 5.

Gerfen CR, Surmeier DJ (2011) Modulation of striatal projection systems by dopamine. Annu Rev Neurosci 34:441–466.

Gertler TS, Chan CS, Surmeier DJ (2008) Dichotomous anatomical properties of adult striatal medium spiny neurons. J Neurosci 28:10814–10824.

Gittis AH, Nelson AB, Thwin MT, Palop JJ, Kreitzer AC (2010) Distinct roles of GABAergic interneurons in the regulation of striatal output pathways. J Neurosci 30:2223–2234.

Goldberg JA, Reynolds JNJ (2011) Spontaneous firing and evoked pauses in the tonically active cholinergic interneurons of the striatum. Neuroscience 198:27–43.

Groenewegen HJ, Berendse HW (1990) Connections of the subthalamic nucleus with ventral striatopallidal parts of the basal ganglia in the rat. J Comp Neurol 294:607–622.

Gundersen HJ, Jensen EB, Kiêu K, Nielsen J null (1999) The efficiency of systematic sampling in stereology--reconsidered. Journal of Microscopy 193:199–211.

Guo Q, Wang D, He X, Feng Q, Lin R, Xu F, Fu L, Luo M (2015) Whole-brain mapping of inputs to projection neurons and cholinergic interneurons in the dorsal striatum. PLoS ONE 10:e0123381.

Ibáñez-Sandoval O, Tecuapetla F, Unal B, Shah F, Koós T, Tepper JM (2011) A novel functionally distinct subtype of striatal neuropeptide Y interneuron. The Journal of Neuroscience: The Official Journal of the Society for Neuroscience 31:16757–16769.

Kita H, Kitai ST (1987) Efferent projections of the subthalamic nucleus in the rat: light and electron microscopic analysis with the PHA-L method. J Comp Neurol 260:435–452.

Klug JR, Engelhardt MD, Cadman CN, Li H, Smith JB, Ayala S, Williams EW, Hoffman H, Jin X (2018) Differential inputs to striatal cholinergic and parvalbumin interneurons imply functional distinctions. Elife 7.

Koós T, Tepper JM (1999) Inhibitory control of neostriatal projection neurons by GABAergic interneurons. Nat Neurosci 2:467–472.

Koós T, Tepper JM, Wilson CJ (2004) Comparison of IPSCs evoked by spiny and fastspiking neurons in the neostriatum. J Neurosci 24:7916–7922.

Koshimizu Y, Fujiyama F, Nakamura KC, Furuta T, Kaneko T (2013) Quantitative analysis of axon bouton distribution of subthalamic nucleus neurons in the rat by single neuron visualization with a viral vector. J Comp Neurol 521:2125–2146.

Lee T, Kaneko T, Taki K, Mizuno N (1997) Preprodynorphin-, preproenkephalin-, and preprotachykinin-expressing neurons in the rat neostriatum: an analysis by immunocytochemistry and retrograde tracing. J Comp Neurol 386:229–244.

Mallet N, Micklem BR, Henny P, Brown MT, Williams C, Bolam JP, Nakamura KC, Magill PJ (2012) Dichotomous organization of the external globus pallidus. Neuron 74:1075–1086.

Mallet N, Pogosyan A, Márton LF, Bolam JP, Brown P, Magill PJ (2008a) Parkinsonian beta oscillations in the external globus pallidus and their relationship with subthalamic nucleus activity. The Journal of Neuroscience: The Official Journal of the Society for Neuroscience 28:14245–14258.

Mallet N, Pogosyan A, Sharott A, Csicsvari J, Bolam JP, Brown P, Magill PJ (2008b) Disrupted dopamine transmission and the emergence of exaggerated beta oscillations in subthalamic nucleus and cerebral cortex. The Journal of Neuroscience: The Official Journal of the Society for Neuroscience 28:4795–4806.

Melendez-Zaidi AE, Lakshminarasimhah H, Surmeier DJ (2019) Cholinergic modulation of striatal nitric oxide-producing interneurons. Eur J Neurosci 50:3713–3731.

Monteiro P, Barak B, Zhou Y, McRae R, Rodrigues D, Wickersham IR, Feng G (2018) Dichotomous parvalbumin interneuron populations in dorsolateral and dorsomedial striatum. J Physiol (Lond) 596:3695–3707.

Nakano K, Hasegawa Y, Tokushige A, Nakagawa S, Kayahara T, Mizuno N (1990) Topographical projections from the thalamus, subthalamic nucleus and pedunculopontine tegmental nucleus to the striatum in the Japanese monkey, Macaca fuscata. Brain Res 537:54–68.

Nambu A, Tokuno H, Takada M (2002) Functional significance of the cortico-subthalamo-pallidal “hyperdirect” pathway. Neurosci Res 43:111–117.

Pan WX, Mao T, Dudman JT (2010) Inputs to the dorsal striatum of the mouse reflect the parallel circuit architecture of the forebrain. Frontiers in Neuroanatomy 4:147.

Planert H, Szydlowski SN, Hjorth JJJ, Grillner S, Silberberg G (2010) Dynamics of synaptic transmission between fast-spiking interneurons and striatal projection neurons of the direct and indirect pathways. J Neurosci 30:3499–3507.

Rico AJ, Barroso-Chinea P, Conte-Perales L, Roda E, Gómez-Bautista V, Gendive M, Obeso JA, Lanciego JL (2010) A direct projection from the subthalamic nucleus to the ventral thalamus in monkeys. Neurobiol Dis 39:381–392.

Sato F, Parent M, Levesque M, Parent A (2000) Axonal branching pattern of neurons of the subthalamic nucleus in primates. The Journal of Comparative Neurology 424:142–152.

Schweizer N, Viereckel T, Smith-Anttila CJA, Nordenankar K, Arvidsson E, Mahmoudi S, Zampera A, Wärner Jonsson H, Bergquist J, Lévesque D, Konradsson-Geuken Å, Andersson M, Dumas S, Wallén-Mackenzie Å (2016) Reduced Vglut2/Slc17a6 Gene Expression Levels throughout the Mouse Subthalamic Nucleus Cause Cell Loss and Structural Disorganization Followed by Increased Motor Activity and Decreased Sugar Consumption. eNeuro 3:ENEURO.0264-16.2016.

Sharott A, Vinciati F, Nakamura KC, Magill PJ (2017) A Population of Indirect Pathway Striatal Projection Neurons Is Selectively Entrained to Parkinsonian Beta Oscillations. J Neurosci 37:9977–9998.

Silberberg G, Bolam JP (2015) Local and afferent synaptic pathways in the striatal microcircuitry. Curr Opin Neurobiol 33:182–187.

Smith JB, Klug JR, Ross DL, Howard CD, Hollon NG, Ko VI, Hoffman H, Callaway EM, Gerfen CR, Jin X (2016) Genetic-Based Dissection Unveils the Inputs and Outputs of Striatal Patch and Matrix Compartments. Neuron 91:1069–1084.

Smith Y, Bevan MD, Shink E, Bolam JP (1998) Microcircuitry of the direct and indirect pathways of the basal ganglia. Neuroscience 86:353–387.

Smith Y, Hazrati LN, Parent A (1990) Efferent projections of the subthalamic nucleus in the squirrel monkey as studied by the PHA-L anterograde tracing method. J Comp Neurol 294:306–323.

Smith Y, Parent A (1988) Neurons of the subthalamic nucleus in primates display glutamate but not GABA immunoreactivity. Brain Res 453:353–356.

Szydlowski SN, Pollak Dorocic I, Planert H, Carlén M, Meletis K, Silberberg G (2013) Target selectivity of feedforward inhibition by striatal fast-spiking interneurons. The Journal of Neuroscience: The Official Journal of the Society for Neuroscience 33:1678–1683.

Tepper JM, Koós T, Ibanez-Sandoval O, Tecuapetla F, Faust TW, Assous M (2018) Heterogeneity and Diversity of Striatal GABAergic Interneurons: Update 2018. Frontiers in Neuroanatomy 12:91.

Tepper JM, Koós T, Wilson CJ (2004) GABAergic microcircuits in the neostriatum. Trends Neurosci 27:662–669.

Tepper JM, Tecuapetla F, Koós T, Ibáñez-Sandoval O (2010) Heterogeneity and diversity of striatal GABAergic interneurons. Frontiers in Neuroanatomy 4:150.

Tervo DGR, Hwang B-Y, Viswanathan S, Gaj T, Lavzin M, Ritola KD, Lindo S, Michael S, Kuleshova E, Ojala D, Huang C-C, Gerfen CR, Schiller J, Dudman JT, Hantman AW, Looger LL, Schaffer DV, Karpova AY (2016) A Designer AAV Variant Permits Efficient Retrograde Access to Projection Neurons. Neuron 92:372–382.

Wall NR, De La Parra M, Callaway EM, Kreitzer AC (2013) Differential innervation of direct-and indirect-pathway striatal projection neurons. Neuron 79:347–360.

Wallén-Mackenzie Å, Dumas S, Papathanou M, Martis Thiele MM, Vlcek B, König N, Björklund ÅK (2020) Spatio-molecular domains identified in the mouse subthalamic nucleus and neighboring glutamatergic and GABAergic brain structures. Commun Biol 3:338.

West MJ, Slomianka L, Gundersen HJ (1991) Unbiased stereological estimation of the total number of neurons in thesubdivisions of the rat hippocampus using the optical fractionator. Anat Rec 231:482–497.

